# Reverse transcriptase inhibitors induce autophagy in a LINE-1 ORF1p-dependent manner

**DOI:** 10.1101/2025.08.07.669170

**Authors:** Michela Damizia, Mirko Baranzini, Ilaria Sciamanna, Ludovica Altieri, Paola Rovella, Omar G. Rosas Bringa, Samira Hozeifi, Leila J Saba, Apostolos Mourtzinos, Anna O’Shea, Mohammad Salik Zeya Ansari, Federica Andreola, Roberto Cirilli, Gianluca Sbardella, Annalucia Serafino, Gerald G. Schumann, Daniela Trisciuoglio, John LaCava, Patrizia Lavia, Corrado Spadafora

## Abstract

Human <u>L</u>ong Interspersed <u>N</u>uclear <u>E</u>lement-1 (LINE-1) retrotransposons propagate throughout the genome via reverse-transcribed RNA intermediates. LINE-1 expression is pervasive in cancer. Functional LINE-1s encode two proteins: ORF1p, an RNA-binding protein, and ORF2p, harboring reverse transcriptase and endonuclease activities. Reverse transcriptase inhibitors, including non-nucleoside (NNRTI) and nucleoside (NRTI) inhibitors, inhibit cancer cell proliferation and antagonize cancer progression. We previously found that two NNRTIs induced DNA damage, nuclear lamin rupture, micronuclei formation, and autophagy in prostate cancer cells. We now find that two different RTIs up-regulate LINE-1 mRNA expression and ORF1p abundance in nuclei, triggering ORF1p interactions with lamin B1 and with DNA damage factors. ORF1p accumulates within micronuclei with damaged DNA and with the autophagy receptor p62. We further demonstrate that inhibiting autophagy, or decreasing ORF1p levels, prevent DNA damage and preserve lamin B1 integrity, uncoverig a role of LINE-1-ORF1p in the autophagy response of cancer cells, independent on retrotranscription events.

## Introduction

LINE-1/L1 retrotransposons comprise the only autonomous retrotransposon family in humans. The human haploid genome contains ∼500,000 LINE-1 copies, accounting for about 18% of the genomic DNA content^1^. The vast majority of LINE-1 copies are not retrotransposition-competent, being 5’- truncated, or internally rearranged or harbor mutations, yet a subset of ∼80-100 LINE-1 elements is intact and retrotransposition-competent^2, 3^. Intact LINE-1 elements (∼6 kb in length), harboring a 5’- and a 3’- untranslated region (UTR), and end in a poly(A) tail (^4^ and references herein). The resulting full-length mRNA is bicistronic and harbors two open reading frames separated by a 63-nt intergenic region: ORF1 encodes ORF1p, a ∼40 kDa RNA-binding protein, and ORF2 codes for ORF2p, a 150 kDa polyprotein with reverse transcriptase (RT) and endonuclease (EN) activities^4^. ORF1p and ORF2p are both essential for LINE-1 retrotransposition through “target-primed reverse transcription”^5, 6^ of an mRNA intermediate that is reverse-transcribed into cDNA at the genomic integration site of the new LINE-1 copy. In the process, the ORF2p protein EN and RT domains are responsibe for genomic DNA cleavage, and for reverse transcription of LINE-1 mRNA in a cDNA copy, respectively. ORF1p is a shuttling protein involved in formation and transport of ribonucleocomplexes in and out of the nucleus, but its precise role remains less well understood.

LINE-1 mRNA is highly expressed in pluripotent stem cells, while being barely detectable in differentiated cell types^7^. LINE-1 expression is modulated during differentiation, generates somatic mosaicism during development^8^, and plays roles in de- and recondensation of chromatin in embryonic preimplantation stages^9,10^, indicating that LINE-1 activity is integral to early developmental processes. LINE-1 elements are gradually repressed during differentiation^11^, except in neurons, which remain retrotransposition-permissive (reviewed in^12, 13^): in that context, active LINE-1s provide a source of regulatory RNAs acting in specific developmental windows and/or cell types during brain development and differentiation^14^. Indeed, transdifferentiation of murine embryonic fibroblasts to dopaminergic neurons entails the reactivation of LINE-1, which modulate chromatin opening and transcription^15^ and genome-wide chromatin remodeling in the developing brain^16^.

Full-length LINE-1s are dysregulated in de-differentiated, proliferating cancer cells and exhibit abnormally high expression, with ORF1p emerging as a hallmark of almost half of human cancers^17^. Many tumor types show elevated LINE-1 expression^18^ and accumulation of polymorphic LINE-1 *de novo* insertions in the genome^19,20^; reviewed in ^21–24^. Although most LINE-1 insertions are viewed as “passenger” mutations rather than drivers in cancer^25, 26^; reviewed in^27^, the capacity of LINE-1 to drive extensive, megabase-scale genomic rearrangements is documented across numerous cancer genomes: LINE-1 insertions can orchestrate cancer-promoting conditions by inactivating regions containing tumor suppressor genes or driving chromosomal rearrangements that lead to the amplification of oncogenes^26^. Conversely, reducing LINE-1 expression in RNA interference (RNAi) assays inhibits cancer cell proliferation, associated with a substantial remodulation of the expression profile of small regulatory RNAs – including miRNAs and piRNAs – and ensuing global changes in gene expression^28–30^.

There is evidence that certain RT inhibitors (RTIs) designed to inhibit the HIV-1 RT activity also modulate cell proliferation in non-HIV-infected cancer cells. RTI molecules belong to two main classes: non-nucleoside RT inhibitors (NNRTIs), which bind the HIV-1 RT enzyme and allosterically block its activity, and nucleoside RT inhibitors (NRTIs), which mimic nucleotide substrates in the enzyme active site, inhibiting cDNA elongation. Many NRTIs also inhibit LINE-1-encoded RT. Most NNRTIs are instead considered to be HIV-specific^31^, although efavirenz (EFV) has been reported to reduce LINE-1 retrotransposition efficiency *in vitro*^32^. RTIs of both classes exert widespread effects in human cells. NNRTIs (e.g. EFV, nevirapine, dapivirine, doravirine) reduce cell proliferation and activate apoptosis in cancer cell lines, and inhibit progression of various cancer types in animal models^28; 33–45^. EFV also showed therapeutic efficacy in a phase-II clinical trial with metastatic castration-resistant prostate cancer patients^46,47^. A new generation structure-activity relationship (SAR)-designed compound, SPV122, a member of pyrimidinone-derived NNRTIs^48,49^, also exhibited anticancer activity in cell lines and animal models^50,51^, particularly the purified SPV122.2 stereoisomer. NRTIs (e.g. Abacavir, ABC; Azidothymidine, AZT; Lamivudine) have similar antiproliferative effects^43, 45, 52–53^.

In all studied cases, discontinuation of the RTI treatment led to resumption of cancer progression^28, 29, 51^: this suggests that the cancer inhibitory effects observed in cell lines, animal models and human trials are independent of LINE-1 *de novo* insertions that would arise from LINE-1 retrotransposition. Indeed, we previously found that RTI administration to cancer cells – similar to LINE-1 RNAi – impairs the biogenesis of miRNAs with pro-oncogenic roles and reprograms miRNA transcription, restoring a global gene expression profile compatible with the non-cancer state^38^. Furthermore, EFV and SPV122.2 induced genome damage, methylated H3 histone reorganization, lamin B1 fragmentation and the appearance of micronuclei, concomitant with autophagy activation and cell death induction in prostate cancer cells^54^, and similar effects were observed in BRAF-mutant melanoma cells^55^. NNRTIs triggered these responses in cancer cells, but not in non-cancer cell lines, implying a selective vulnerability of cancer cells.

Here we have sought to clarify whether LINE-1 mRNA and proteins, which are overexpressed in cancer cells, play roles in modulating cellular changes induced by RTIs. We report that, unexpectedly, full-length LINE-1 mRNA expression and ORF1p abundance increase in prostate cancer cells upon exposure to both NNRTIs (EFV and SPV122.2), and to an NRTI (abacavir, ABC), acting as stressors. Following that increase, RTIs of both classes induce novel interactions of ORF1p with DNA damage factors and with lamin B1. We find that ORF1p concentrates, together with the damaged DNA, in micronuclei formed during RTI treatment, which furher recruit the autophagy receptor p62. Moreover, LINE-1 expression and autophagy activation are both required for RTI induction of genome damage and nuclear lamina reuptures. These data highlight a role of LINE-1 and its product ORF1p, which is clearly uncoupled from reverse transcription-dependent events and involves autophagy effectors that ultimately control cell fates.

## Results

### The NNRTIs EFV and SPV122.2 and the NRTI ABC increase L1 mRNA transcription and ORF1p protein abundance

Molecules of both the NNRTI and NRTI classes, albeit being designed to inhibit HIV RT, affect multiple functions in HIV non-infected cells, as summarized in the introduction. In particular, the metastatic prostate cancer cell line PC3 was characterized in previous studies and proved sensitive to cell proliferation inhibition when cultured with NNRTIs, while non cancer prostate epithelial cells were minimally affected, if at all^54^. In order to assess whether these RTIs affect endogenous LINE-1 retrotransposon expression, we analyzed LINE-1 mRNA abundance in PC3 cells cultured in the presence and absence of the NNRTIs EFV and SPV122.2, and of the NRTI ABC – the latter has established inhibitory capacity over LINE-1-encoded RT activity^31^. PC3 cell cultures were treated with RT inhibitors (2 cycles of 96 h each, see Methods), a schedule that effectively reduces cell proliferation^54^; control cultures were either treated with DMSO only (control for EFV and SPV122.2, both of which were dissolved in DMSO), or untreated (control for treatment with the water-soluble ABC). All samples were analyzed for endogenous LINE-1 mRNA levels using primers specific for *ORF1* and *ORF2* coding regions (map Fig. 1A) in q-RT-PCR assays. Unexpectedly, all three RTIs induced a statistically significant increase in LINE-1 mRNA levels in PC3 cells compared to controls (1.3- to 2.6-fold, Fig. 1B), with SPV122.2 being most effective. We asked whether the increase in LINE-1 mRNA levels was accompanied by a corresponding increase in abundance of L1-ORF1p. Immunoblotting of PC3 cell extracts revealed a statistically significant increase in the overall amount of ORF1p (1.6 to 1.7-fold) in PC3 cultures treated with all three RTIs compared to controls (Fig.1C). ORF1p was further analyzed by automated quantitative immunofluorescence (IF; exemplifying fields are shown in Fig. 1D-E). The analysis of the distribution of ORF1p fluorescence signals in single cells revealed a highly significant increase in PC3 cultures treated with both SPV122.2 and ABC compared to their controls (Fig. 1D-E). ORF1p is an mRNA chaperone and packaging protein that is predominantly located in the cytoplasm^56–58^. We analyzed the subcellular distribution of ORF1p signals by automated fluorescence, selecting either whole cells or the DAPI-stained mask depicting nuclei as the Region of Interest (RoI; exemplified in Fig. 1D). PC3 control cultures (DMSO, CTR) showed faint ORF1p fluorescence signals, largely confined to the cytoplasm. Their intensity increased significantly in RTI-treated cells, particularly in nuclei (Fig. 1D-E). These data indicate that both SPV122.2 and ABC up-regulate LINE-1 expression at both RNA and ORF1p levels and that this increase leads to an increased accumulation of L1-ORF1p in the nuclei of PC3 prostate cancer cells.

**Fig. 1.**
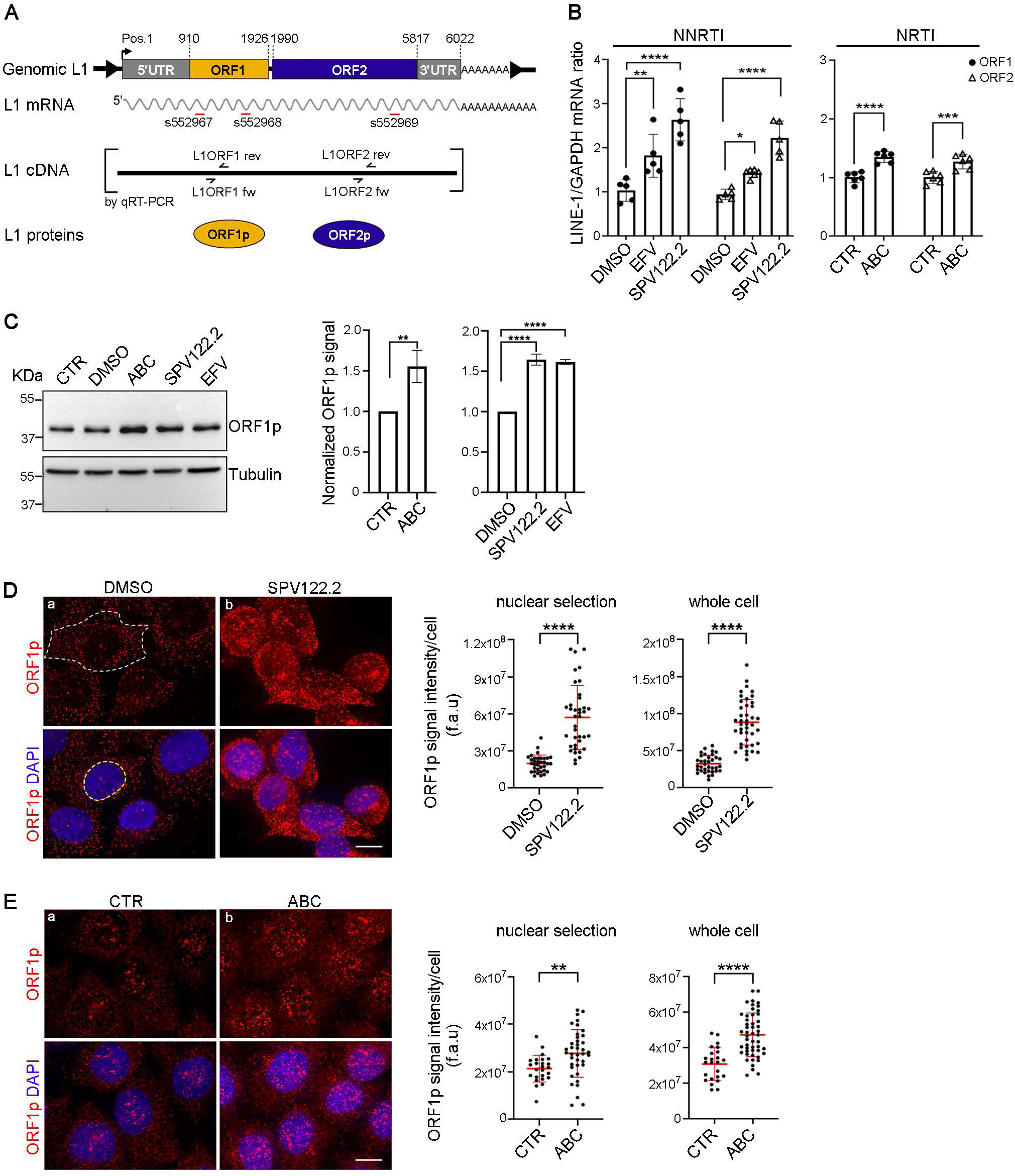
SPV122.2 and EFV (NNRTI class), and Abacavir (NRTI class) up-regulate LINE-1 mRNA and ORF1p levels in PC3 cells. **A.** Structure of a functional human L1 Hs retrotransposon. The schematic shows the binding sites for qPCR primers (L1ORF1 For, L1 ORF1 Rev, L1ORF2 For and L1ORF2 Rev), represented as small convergent black arrows on L1 cDNA, and for siRNAs s552967, s552968 and s552969 (red lines; coordinates in the Methods section) on full-length L1 mRNA. **B.** RT-PCR quantification of endogenous full-length L1 mRNA transcripts using *ORF1*- and *ORF2*-specific primers shown in **A** and RNA extracted from PC3 cells cultured with EFV, SPV122.2 or ABC (2 cycles, 96 h each) relative to their respective controls (controls are set as 1). Throughout this study, DMSO-soluble NNRTIs were compared to cultures treated with 0.2% DMSO alone (indicated as DMSO), while the hydrosoluble ABC was compared to untreated control cultures (indicated as CTR). Data are presented as the mean ± SD from 8 assays. Statistical significance was assessed using the two-way ANOVA and Multiple Comparison Test. **, p<0.01; ***, p<0.001; ****, p<0.0001. **C.** Representative immunoblot assay of ORF1p in PC3 whole cell extracts cultured with and without SPV122.2, EFV or ABC; tubulin was used as a loading control. In three independent immunoblotting assays, ORF1p signals were measured by densitometry and normalized to those of the loading control (either GAPDH, or tubulin, or both were used in different experiments). Histograms represent the mean ± SD ratios, setting the values obtained in untreated and in DMSO-treated cells as 1. Statistical significance was calculated using the unpaired Student’s *t-*test, with **, p<0.01 (ABC vs. CTR) and ****, p<0.0001 (SPV122.2 vs DMSO and EFV vs. DMSO). **D-E.** Immunofluorescence (IF) fields immunostained for ORF1p in SPV122.2 vs. DMSO-treated **(D),** and ABC-treated vs. untreated (CTR) **(E)** PC3 cultures. Bars: 10 μm. The scatter plots on the right, display the distribution of ORF1p signal intensity per cell measured in the experiments in **D** and **E** (30-50 counted cells per sample). The ORF1p fluorescence intensity was measured both in whole cells (in **D,** panel a, the white dashed line delimits one exemplifying cell profile), and after selecting DAPI-stained nuclei (the yellow dashed line in **D,** panel b, delimits one exemplifying nucleus) as the Region of Interest (RoI). The distributions of signals were statistically analyzed using the Mann-Whitney test: **, p<0.01; ***, p<0.001; ****, p<0.0001. The same trend was replicated in three independent experiments for SPV122.2 (140 DMSO and 120 SPV122.2 analyzed cells), and in four independent experiments for ABC (180 CTR and 250 ABC analyzed cells).

### Both ABC and SPV122.2 induce nuclear accumulation of the DNA damage marker γ−H2AX and of the stress-responsive factor BCLAF1 in association with L1-ORF1p

We previously showed that the capacity of the NNRTIs EFV and SPV122.2 to inhibit PC3 cell proliferation was accompanied by the induction of DNA damage and activation of the DNA damage response (DDR), with the accumulation of the DNA damage marker γ−H2AX at double-strand breaks (DSB) and CHK2 kinase activation^54^. Those results were consistent with earlier findings indicating that increased L1 expression yielded DSB formation^59^. Concurrently, EFV and SPV122.2 activated autophagy^54^. Given that the NRTI ABC also induces an increase in L1-ORF1p levels (Fig. 1C-D), we wondered whether it similarly triggers an autophagic response. To address that question, we compared the distribution of LC3-II, a marker of autophagosome formation, and p62, an early autophagy receptor, by quantitative IF in PC3 cultures treated with SPV122.2 (Fig. S1A) and ABC (Fig. S1B). A significant increase was detected for both autophagy markers in the RTI samples compared to their controls (quantified in the graphs in Fig S1A-B).

To directly address the link between the autophagic response to RTIs (both SPV122.2 and ABC) and DNA damage, we treated PC3 cells according to the treatment schedule in Fig. 2A, with each RTI either alone, or followed by the addition of 3-Methyladenine (3-MA), an early autophagy inhibitor that suppresses autophagosome formation^60^, then analyzed γ-H2AX expression as a reporter of genomic DNA damage. As shown in Fig. 2B, SPV122.2 (panel b) induces the nuclear accumulation of γ−H2AX, as revealed by the increase in fluorescence signal intensities compared to DMSO-treated PC3 cells (panel a), which is consistent with an increase in DSBs formation. 3-MA (panel c) per se does not affect γ−H2AX levels (panel a). However, 3-MA addition (panel c) to SPV122.2- treated cultures counteracts the accumulation of γ−H2AX induced by SPV122.2 (panel d; the statistical analysis is displayed in the graph below). ABC also induces γ−H2AX nuclear accumulation in treated PC3 cells (Fig. 2C, panel b) compared to untreated controls (panel a), which is again abrogated by 3-MA addition (Fig. 2C, panel d and graph below). Thus, the γ−H2AX accumulation induced by both the NRTI and the NNRTI requires the activation of autophagy.

**Fig. 2.**
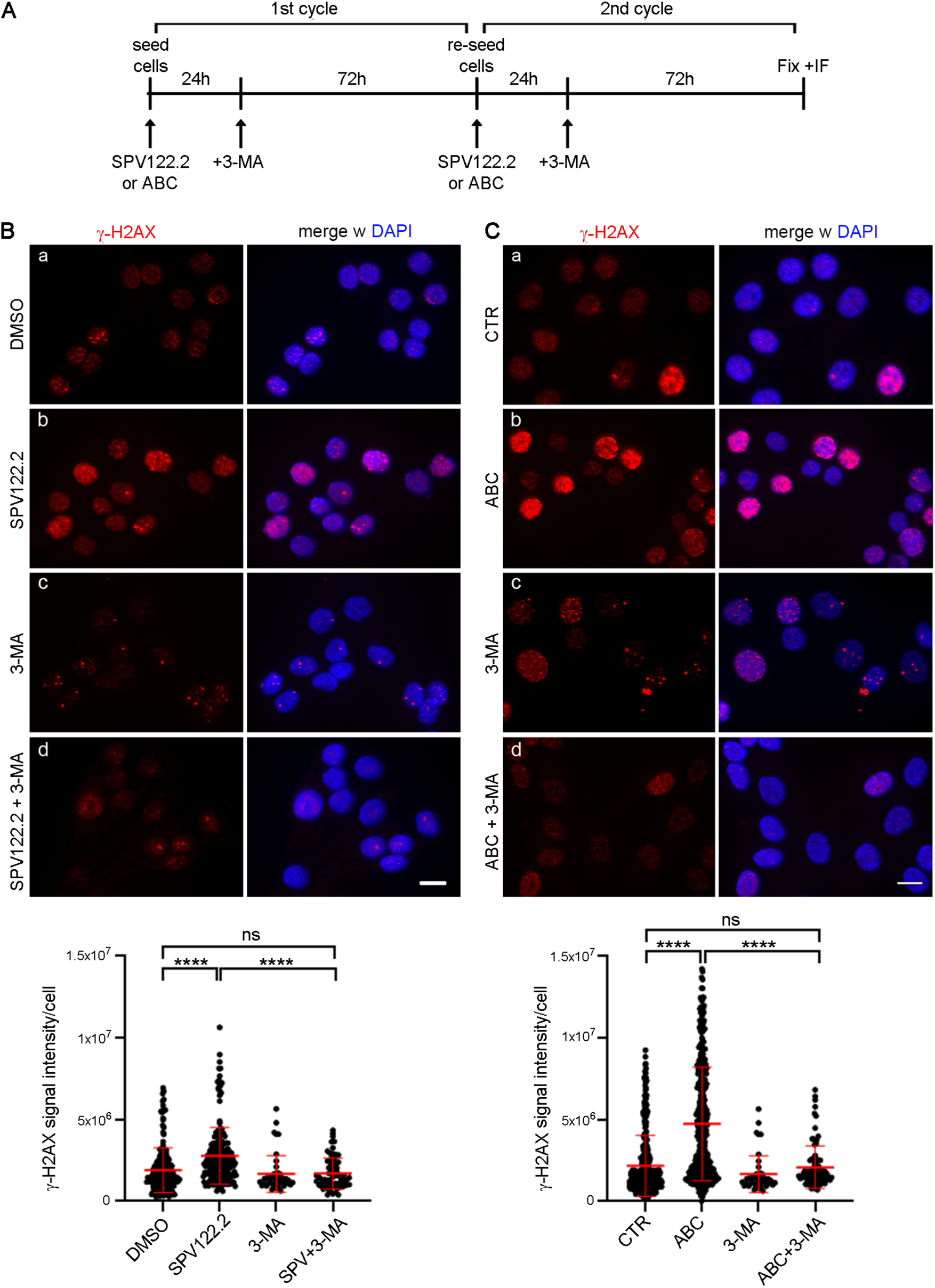
SPV122.2 and ABC inhibitors up-regulate γ−H2AX accumulation in nuclei in an autophagy-dependent manner. **A.** Schedule of treatments used to analyze the implication of autophagy in the induction of DNA damage by RTIs. (3-MA, 3-Methyladenine). **B.** IF analysis of γ−H2AX in SPV122.2-treated PC3 cells: γ−H2AX nuclear signals increased significantly compared to DMSO-treated cultures. Addition of the autophagy inhibitor 3-MA (SPV122.2+3-MA panels) prevents that increase, bringing γ−H2AX signals back to the level measured in DMSO controls. Bar, 10 μm. The graphs below show the distribution of γ−H2AX intensity signals per cell under all tested conditions (>70 counted cells per condition in 2 independent assays; ****, p<0.0001; ns, non significant). **C**. IF analysis of γ−H2AX in ABC-treated PC3 cultures: nuclear γ−H2AX signals significantly increase compared to untreated CTR, and 3-MA addition counteracts that increase. Bar, 10 μm. At least 90 cells per condition were analyzed in 3 independent assays. **, p<0.01; ****, p<0.0001; ns, non significant. The Kruskal-Wallis test (Dunn’s multiple comparison test) was used to statistically analyze the data presented in **B** and **C**.

Since SPV122.2 and ABC increase ORF1p nuclear abundance (Fig. 1C-D) and concomitantly induce DNA damage, we wondered whether the nuclear ORF-1p and γ-H2AX might interact at DSBs. By applying the proximity ligation assay (PLA) methodology, which detects *in situ* interactions or close proximity between endogenous proteins at their subcellular sites, we found that both SPV122.2 (Fig. 3A, panel b) and ABC (Fig. 3B, panel b) increased the formation of nuclear PLA products between ORF1p and γ-H2AX compared to their controls (panels a in Fig. 3A and 3B). These results suggest that both RTIs promote the localization of ORF1p at sites of DNA damage, where ORF1p can interact with γ−H2AX.

**Fig. 3.**
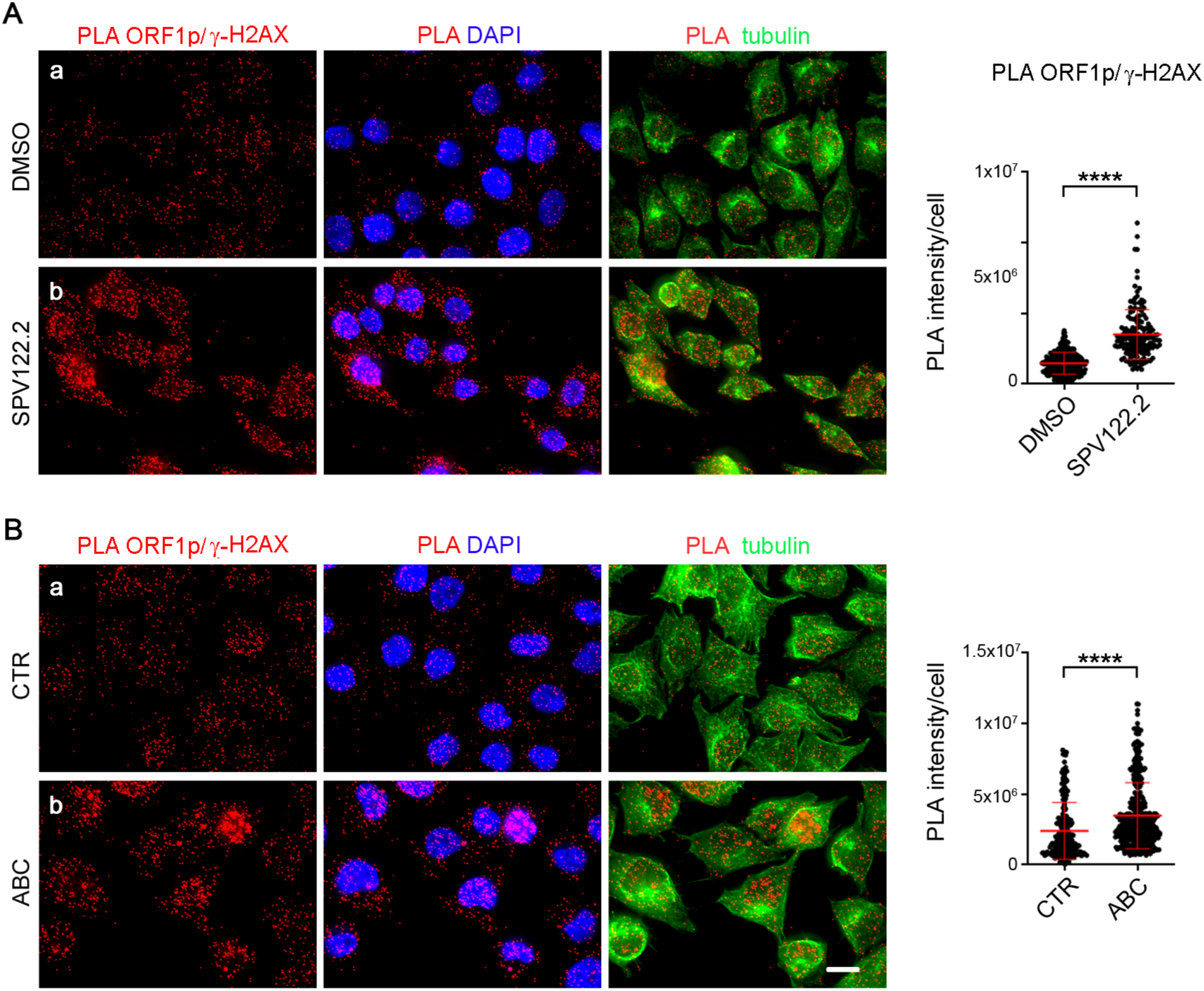
PC3 cell treatment with SPV122.2 and ABC results in an interaction between ORF1p and γ−H2AX. **A.** PLA assays probing ORF1p and γ−H2AX in DMSO- (a) and SPV122.2- (b) treated cultures. **B.** PLA assays between ORF1p and γ−H2AX in untreated CTR (a) and ABC-treated (b) cultures. The PLA probe (red) selectively hybridizes where ligation products are formed. Bars, 10 μm. Treatment with both RTIs up-regulate the formation of ORF1p/γ−H2AX ligation products compared to control cultures. Graphs display the distribution of PLA signals (intensity / cell) measured in 150 to 260 cells per condition in 2 independent assays and statistically analyzed using the Mann-Whitney test. ****, p<0.0001.

Among the factors interacting with γ−H2AX, BCLAF1 is a stress-responsive transcription factor^61^ that localizes predominantly to nuclear speckles^62^ and is recruited to genomic DNA damage sites^63^. BCLAF1 contributes to regulate DNA repair and apoptosis in DNA damaged cells^61^, at least in part via processing of specific RNAs in response to DNA damage^64, 65^. By single cell quantitative IF, BCLAF1 signal intensity significantly increased in both SPV122.2 (Fig. S2A) and ABC (Fig. S2B) - treated cultures compared to their controls. We further analyzed ORF1p/BCLAF1 products in PLA assays. We again detected a significant increase in PLA signals (graphs in Fig. 4) in PC3 samples cultured in the presence of either SPV122.2 (Fig. 4A, panel b) or ABC (Fig. 4B, panel b) compared to their DMSO-treated (Fig. 4A, panel a) or untreated (CTR, Fig. 4B, panel a) controls. Thus the increase in abundance of both γ-H2AX and BCLAF1 induced by both RTIs in consequence of DNA damage is paralleled by an increase in their association with ORF1p.

**Fig. 4.**
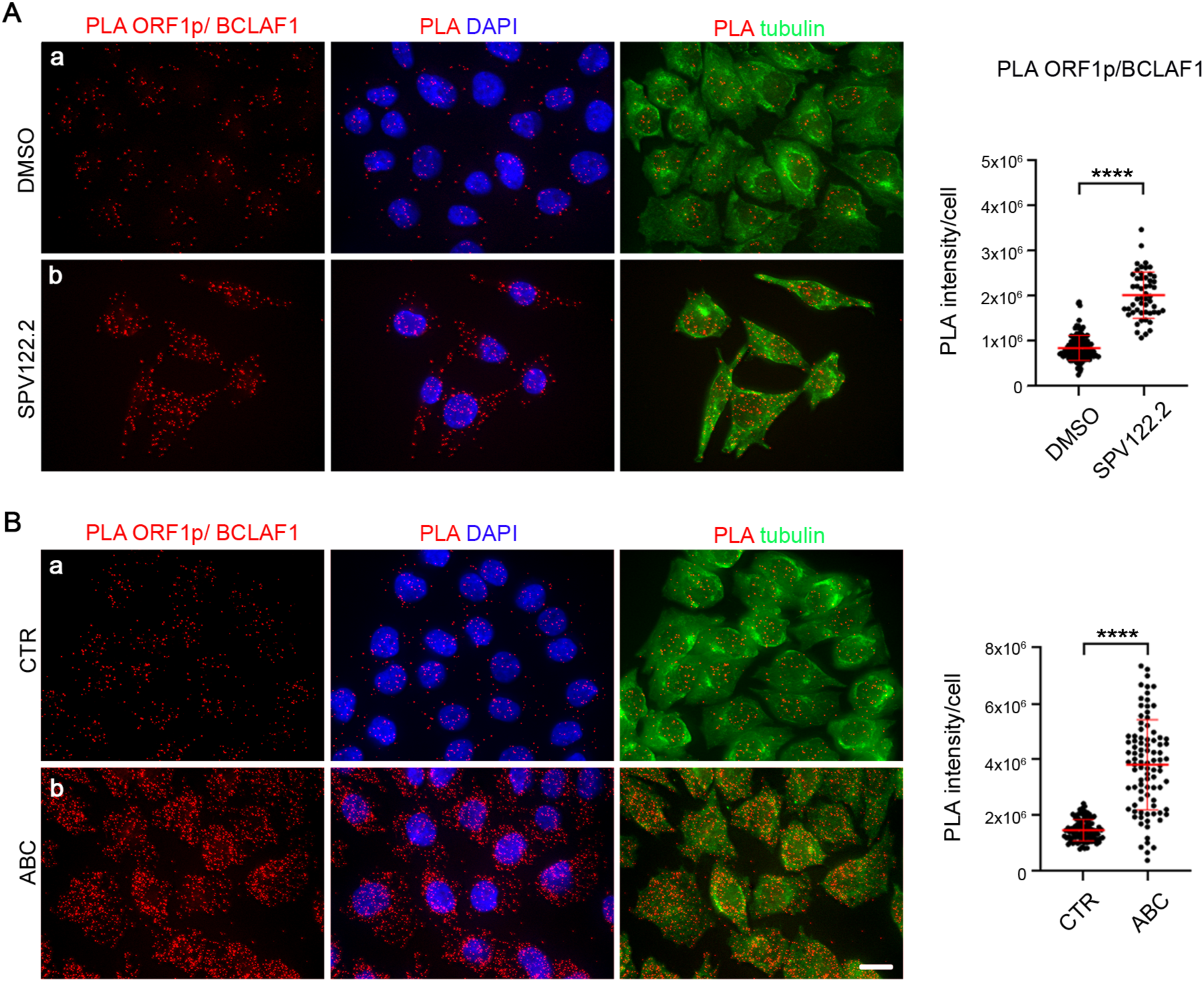
SPV122.2 and ABC induce the formation of ligation products between ORF1p and BCLAF1 proteins in PC3 cells**. A.** PLA assays probing ORF1p and BCLAF1 in DMSO-treated (a) and SPV122.2-treated (b) cultures. **B.** PLA assays probing ORF1p and BCLAF1 in untreated (CTR, a) and ABC-treated (b) PC3 cultures. The PLA probe is depicted in red. Bars, 10 μm. Graphs quantify the PLA signal intensity per cell under each condition, demonstrating an increase in ligation products upon treatment with both RTIs. Cells (>90 cells per sample) were statistically analyzed using the Mann-Whitney test. ****, p<0.0001.

#### Immunoprecipitation-mass spectrometry confirms ORF1p physical interactions

In order to determine if (1) the demonstrated increase in proximal associations between ORF1p and γ-H2AX and between ORF1p and BCLAF1 may extend to other cell models, and if (2) NRTI-induced changes in physical associations are assayable by immunoprecipitation-mass spectrometry (IP-MS), we worked with an alternative cell model and approach. Using the embryonal carcinoma cell line N2102Ep, which exhibits abundant amounts of LINE-1 ORF1p in the steady state^66^, we carried out IP-MS analyses +/- NRTI treatment with stampidine and islatravir for 72 h (see Methods) compared with DMSO carrier as a control. The full list of identified interactors is reported in Table S1. These experiments revealed interactions between ORF1p and γ-H2AX that were enhanced by treatment with either stampidine or islatravir, and interactions between ORF1p and BCLAF1 that were enhanced on treatment with stampidine but not islatravir (Fig. 5). We further highlighted enriched proteins annotated in the Reactome database as belonging to DNA damage and repair pathways and to RNA splicing. Sixteen proteins exhibited increased associations with ORF1p upon treatment with either NRTI used in these assays. The 16 common hits were manually inspected in the Uniprot database (https://www.uniprot.org/). We noticed RNF2, a γ-H2AX-interacting factor that participates in DNA repair^67,68^, and the splicing factor AKAP17A^69^, functionally related to three proteins (MAGOH, RBM25, and SRSF9) annotated in the Reactome RNA splicing pathway. There were also two more hits with roles in steps of the autophagic process: AP2S1-2, a subunit of the endocytic adaptor protein 2 (AP-2) complex^70^, and LAMOR 3^71^. The remaining hits were ribosomal components and factors associated with membranous organelles or vesicles. The interaction of ORF1p with RNA-binding proteins is not surprising given that ORF1p itself is an RNA-binding protein. The interaction with γ-H2AX confirms that the localization of ORF1p at sites of DNA damage is not limited to treatment with SPV122.2 and ABC but can be reproduced using unrelated NRTI molecules.

**Fig. 5.**
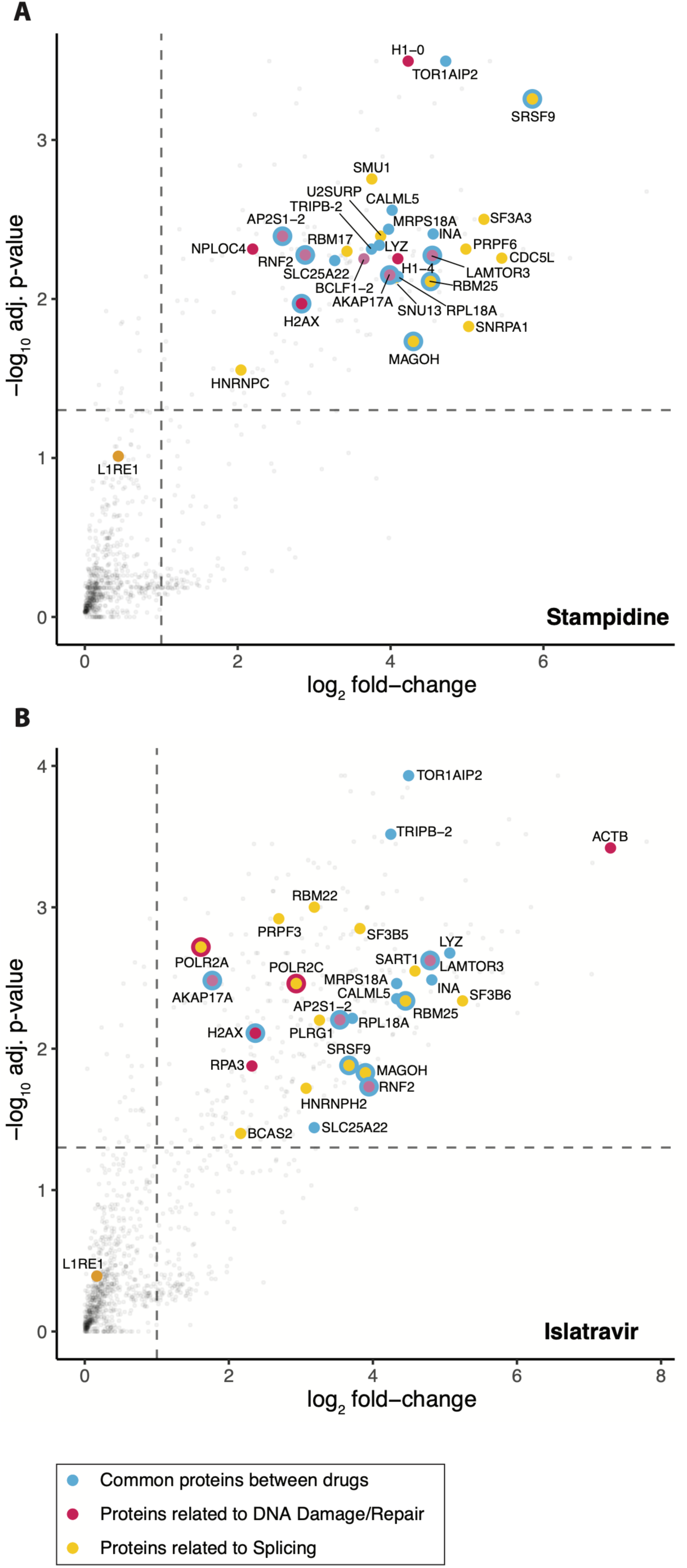
Enrichment of proteins in ⍺-ORF1p IPs after NRTI-treatment of N2102Ep cells. Each scatter plot displays the results of a one-tailed *t*-test for enrichment of proteins associated with ORF1p after NRTI treatment (10 μM in DMSO) of N2102Ep cell cultures, compared against DMSO-treated control cell cultures. **A.** Results from treatment with stampidine. **B.** Results from treatment with islatrivir. Benjamini & Hochberg adjusted -log_10_ p-values are displayed on the y-axis; log_2_ fold-change values are displayed on the x-axis. The horizontal dashed line indicates an adjusted *p*-value = 0.05. The vertical dashed line indicates a fold-change cutoff of 2. Four replicate IPs were used in each treatment and control group. As indicated in the legend: blue dots (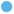) indicate the 16 significantly enriched proteins observed upon both islatravir and stampidine treatments; red dots (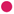) indicate enriched proteins related to DNA damage/repair, according to the Reactome pathway database (identifiers R-HSA-2559586, release V91, and R-HSA-73894, release V91, combined, access January 9, 2025); yellow dots (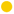) indicate enriched proteins related to splicing, according to Reactome (identifier R-HSA-72203, release V91, access January 13, 2025). Manually selected proteins (BCLAF1-2 and RNF2, DNA repair; AKAP17A, RNA splicing; AP2S1-2 and LAMOR 3, autophagy) are highlighted in magenta (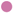), and L1RE1 (ORF1p, the IP target) is highlighted in orange (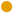). Proteins that meet multiple criteria are indicated by multi-color dots. A complete list of proteins quantified in these analyses is provided in Table S1.

#### SPV122.2 and ABC induce lamin B1 fragmentation and formation of micronuclei that contain ORF1p and p62

Induction of DNA damage can concomitantly cause ruptures in the nuclear envelope (NE) and facilitate the collapse of the nuclear lamina^72, 73^. Lamin B1 is a major nuclear lamina component with critical functions in nuclear architecture and integrity (reviewed in^74^), in chromatin architecture and dynamics^75,76^, and in the response to DNA damage by interacting with the DNA repair factor 53BP1^77^. We previously reported that NNRTIs EFV and SPV122, in addition to inducing DNA damage, also disrupt the nuclear lamina^54^. Fig. S3A shows assays that extend that finding to ABC, which was also found to induce the appearance of nuclei with ruptured lamina that were absent from control PC3 cultures (compare lamin B1 staining in panels a and b), similar to SPV122.2 (Fig. S3B). IF analysis of cells treated with both RTIs showed that lamin B1 fragmentation was associated with increased heterochromatinization, revealed by a boosted signal intensity of dimethyl-H3 histone at lysin 9, H3K9me2 (Fig. S3A-B). Chromatin fragments containing H3K9me2 and associated with lamin B1 fragments were often visualized outside of the nucleus (Fig. S3, arrowed in the zoom-in panels), suggesting the formation of micronuclei in RTI-treated PC3 cultures.

Recent experiments using aneuploidy-inducing anti-mitotic drugs have shown that mis-segregating chromosomes are sensed as DNA damage products and form micronuclei that attract the p62 autophagy receptor^78^. We decided to characterize the micronuclei observed in cells treated with SPV122.2 and ABC in more detail: both drugs increased the formation of micronuclei by 5.5-7-fold (Fig. 6A). We found that these micronuclei contained damaged DNA, revealed by the concentration of γ−H2AX signals, and recruited p62 (Fig. 6B), which is normally a cytoplasmic receptor and is excluded from the nucleus. Furthermore, the micronuclei also recruited ORF1p, which is consistent with the co-localization of ORF1p at sites of damaged DNA detected in PLA assays (Fig. 6C). In these assays, we noticed that p62 accumulated at sites where lamin B1 was heavily fragmented (Fig. 7A): that observation is interesting, as p62 is reported to inhibit the nuclear envelope “sealing” activity of the endosomal sorting complex required for transport III (ESCRT-III), which therefore cannot restore the nuclear envelope integrity^78,79^. ORF1p also localized at sites where the lamin B1 layer exhibited interruptions (Fig. 7B, SPVB122.2, panel b), accumulating particularly where the lamin B1 framework was heavily damaged (Fig. 7B, SPV122.2, panel c, and ABC, panel d). Thus both SPV122.2 and ABC induce ORF1p accumulation at sites of nuclear envelope ruptures and within micronuclei together with γ-H2AX and p62. We wondered whether SPV122.2 and ABC promote the association of ORF1p with lamin B1 itself in PLA assays, as observed for γ−H2AX and BCLAF1. In the absence of SPV122.2 or ABC, we detected only rare, faint ORF1p/lamin B1 PLA signals at the nuclear rim (panels a in Fig. 8A and 8B). Instead, cultures treated with SPV122.2 (Fig. 8A, panel b) or with ABC (Fig. 8B, panel b) displayed abundant PLA products which mainly concentrated at sites neighboring nuclear envelope ruptures (framed cells, zoom-in).

**Fig. 6.**
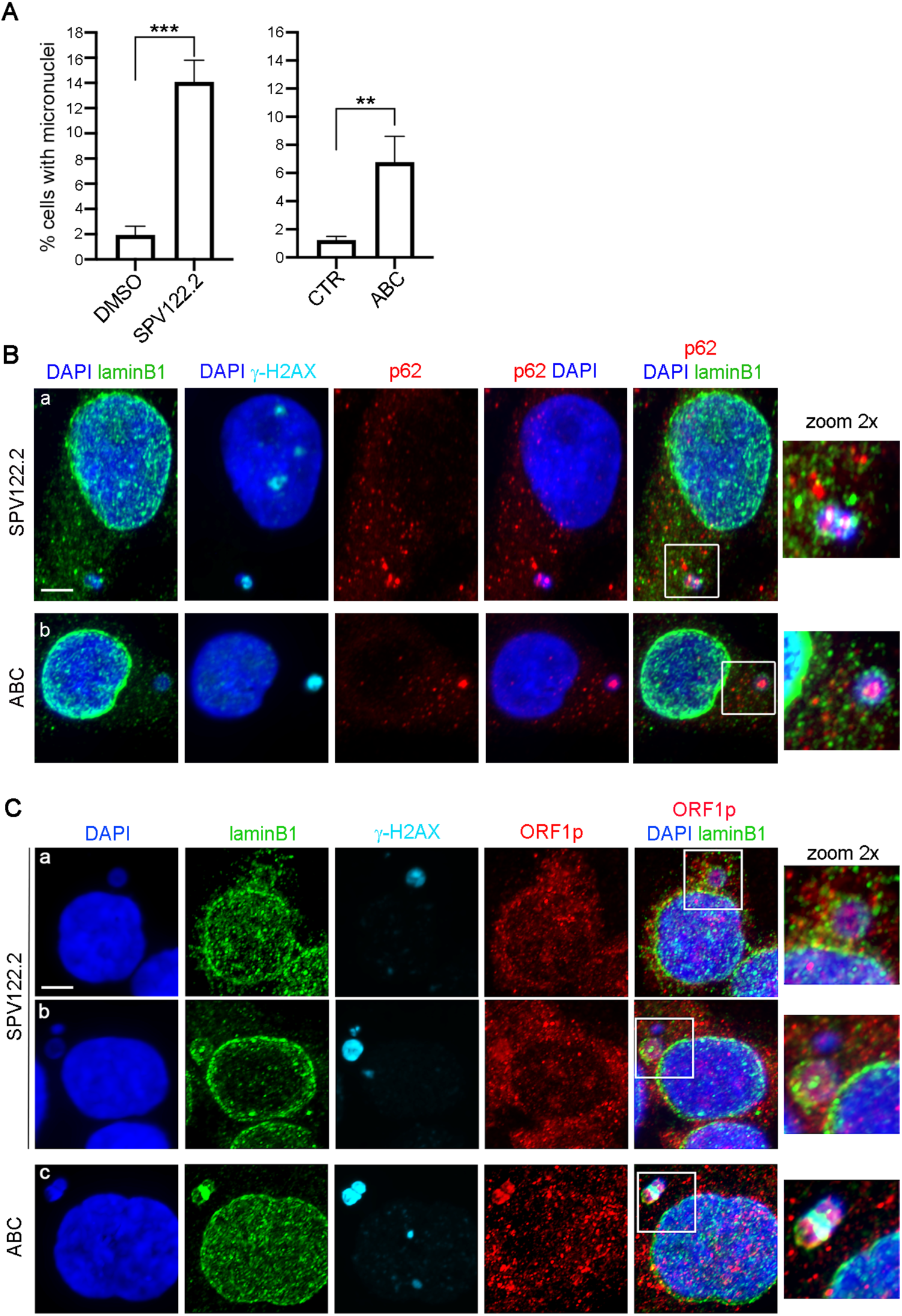
Induction of micronuclei containing ORF1p and p62 in RTI-treated PC3 cultures. **A.** Histograms represent the fraction of cells with micronuclei (mean ± SD) in PC3 cultures treated with RTIs (SPV122.2, ABC), compared to controls (DMSO-treated or untreated, CTR). >450 cells per condition were counted in 3 experiments. SPV122.2-treated were compared to DMSO, and ABC-treated to untreated cultures, using the unpaired *t*-test. **, p <0.01; ***, p <0.001. **B.** Examples of cells stained for lamin B1, γ-H2AX and p62 after treatment with SPV122.2 (panel a) and ABC (panel b): both γ-H2AX and p62 are contained in micronuclei. **C.** Examples of parallel cultures treated with SPV122.2 (panels a-b: two examples are shown) or with ABC (panel c), immunostained for γ-H2AX and ORF1p: both proteins are contained in micronuclei. Micronuclei in **B** and **C** are membrane-enclosed, as depicted by lamin B1 staining.

**Fig. 7.**
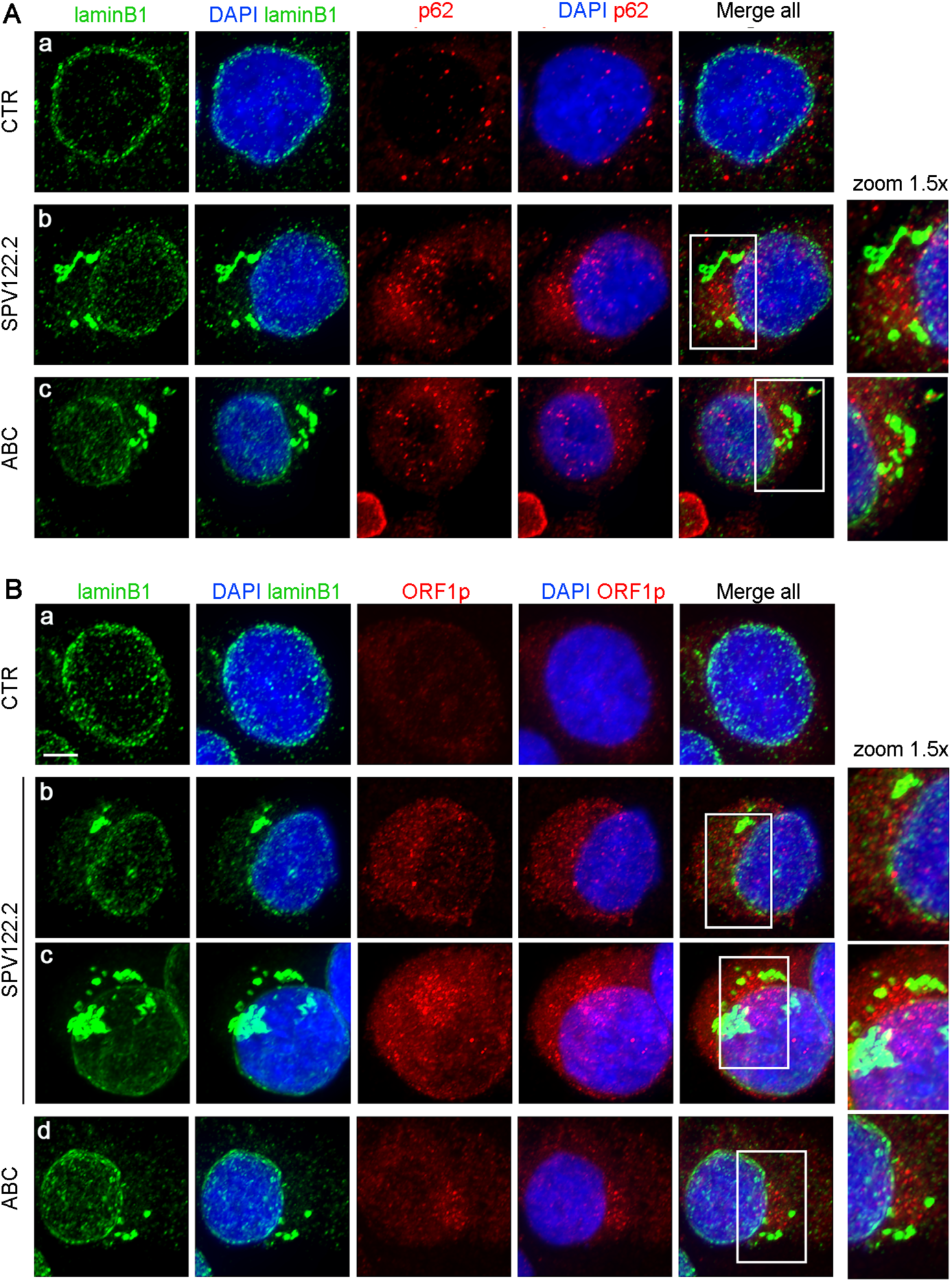
p62 and ORF1p accumulate at sites of lamin B1 ruptures induced by SPV122.2 and ABC**. A.** Exemplifying IF fields from PC3 cultures untreated (a), or treated with SPV122.2 (b) or with ABC (c), immunostained for p62 and lamin B1: p62 accumulates at sites where lamin B1 is fragmented. **B.** Parallel cultures were immunostained for ORF1p and lamin B1: in panel b (SPV122.2), ORF1p signals interpose between discontinuous signals caused by fragmented lamin B1, and result in a punctuated pattern; panels c (SPV122.2) and d (ABC) show examples of more extensive lamin B1 fragmentation, with intense ORF1p accumulation.

**Fig. 8.**
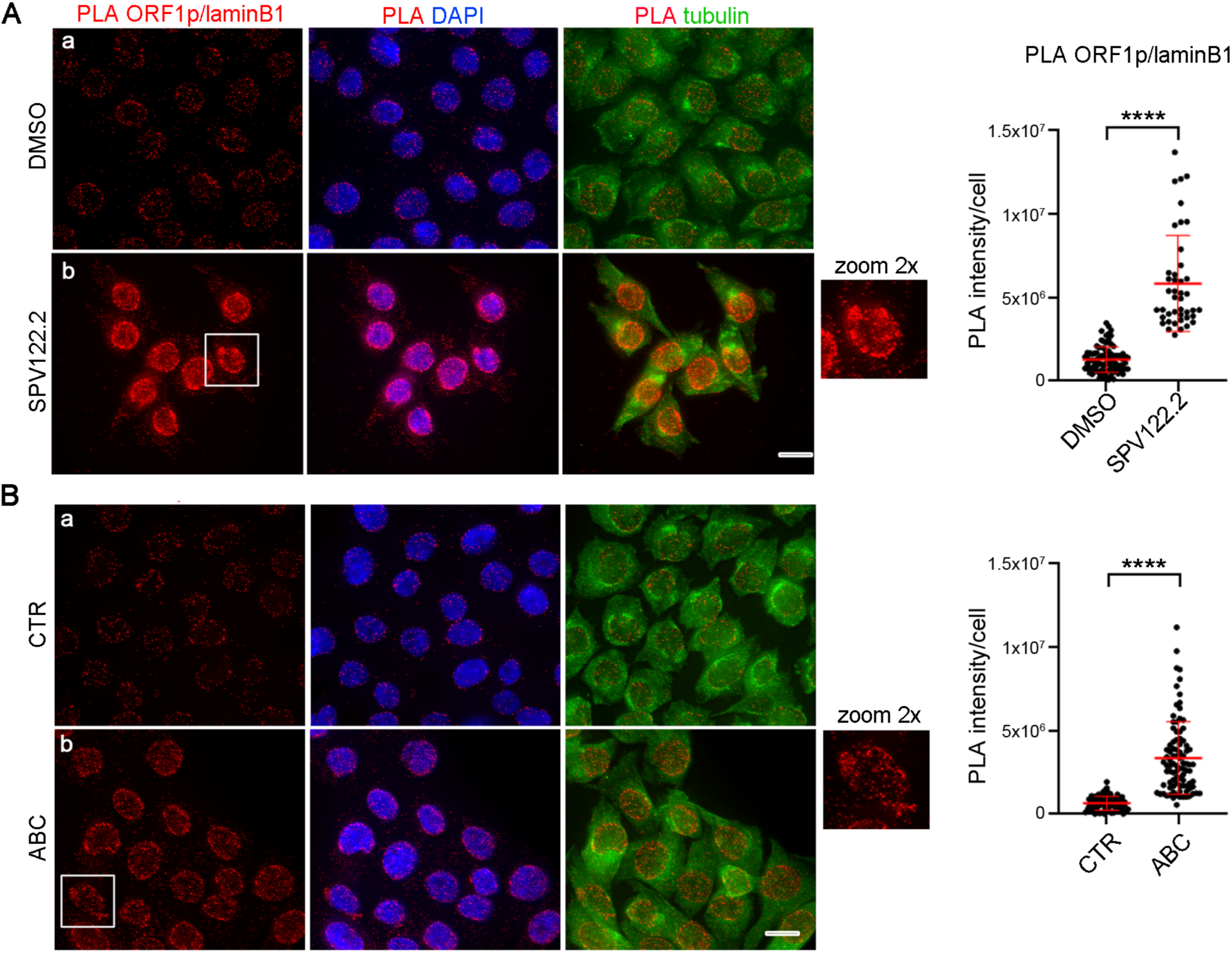
SPV122.2 and ABC induce the formation of ligation products between ORF1p and lamin B1. **A.** PLA assays show that SPV122.2 treatment (panel b) up-regulates ORF1p/lamin B1 ligation products compared to PC3 cultures lacking SPV122 (DMSO-treated, panel a). **B.** PLA assays as in **A** depict a similar increase in ABC-treated (b) compared to untreated (CTR, a) PC3 cultures. In **A** and **B**, red signals locate ligation products to the vicinity of the nuclear envelope, including at sites where the nuclear envelope shows ruptures (framed, 2x zoom-in). Bars, 10 μm. Adjacent graphs show the distribution of PLA signal intensities per cell in presence and absence of SPV122.2 (A) or ABC (B), respectively (>100 analyzed cells per condition). Statistical analysis was performed using the Mann-Whitney test. ****, p<0.0001

#### Autophagy inhibition and interference to LINE-1 expression both prevent lamin B1 fragmentation induced by SPV122.2 and ABC

At this point, we hypothesized that the fragmentation of the nuclear lamina caused by both tested RTIs might be autophagy-related. To assess this possibility, we compared the status of lamin B1 in PC3 cells treated with SPV122.2, either alone or in combination with the autophagy inhibitor 3-MA. The results in Fig. 9A show that 3-MA addition counteracted the disruptive effect of SPV122.2 and preserved the lamin B1 integrity. These findings implicate autophagy in the induction of damage not only at the genome level (as shown in Fig. 2), but also at the level of the nuclear lamina in response to SPV122.2 treatment.

**Fig. 9.**
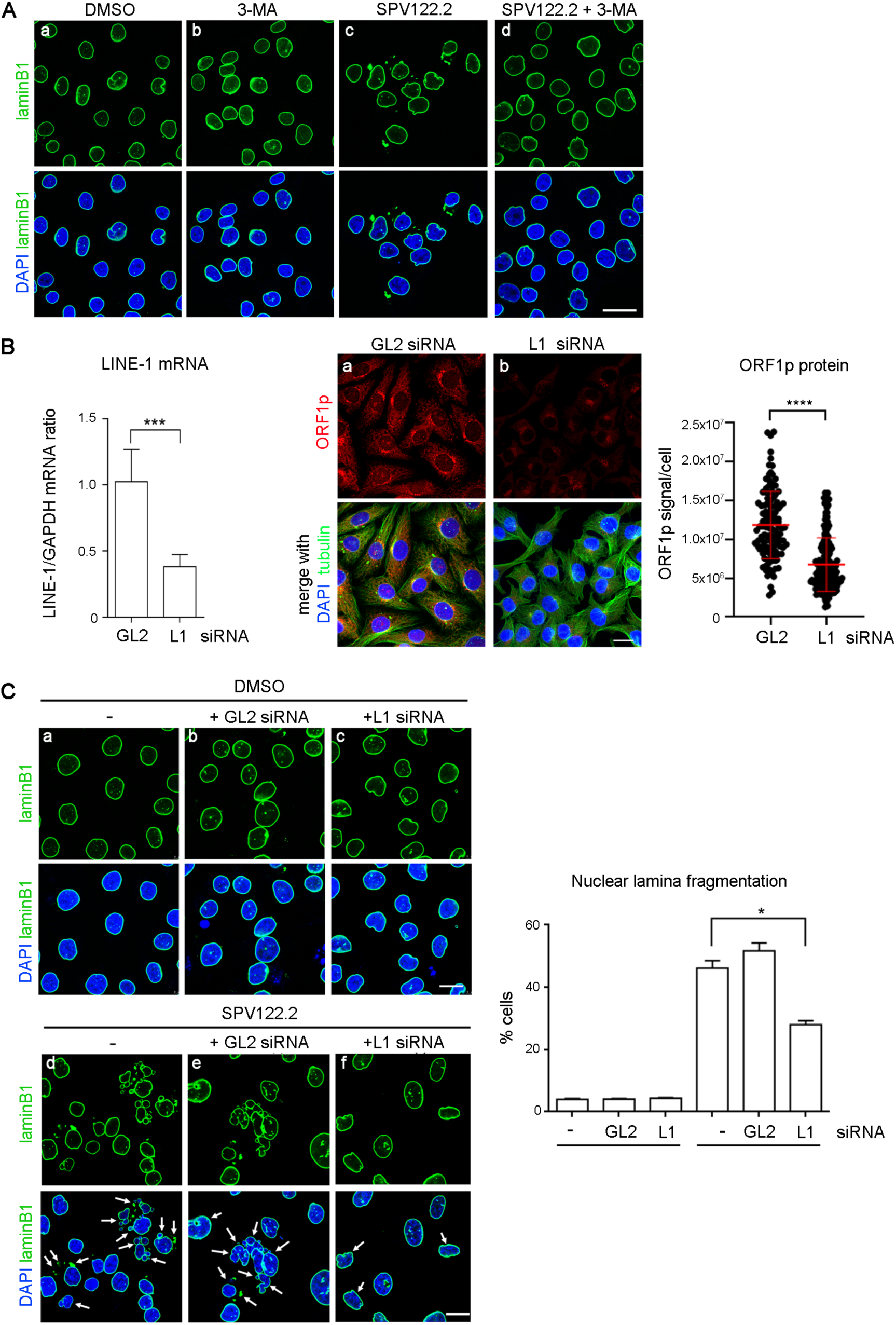
Both autophagy and LINE-1 expression are required for the induction of lamin B1 fragmentation by SPV122.2 and ABC. **A.** Confocal microscopy images of lamin B1 in PC3 cultures: intact nuclear envelopes are depicted by the continuity of lamin B1 staining in DMSO-treated (a) and 3-MA-treated (b) cells; SPV122.2 (c) induces lamin B1 fragmentation and micronuclei formation; 3- MA addition (d) prevents SPV122.2-induced fragmentation. **B.** RT-PCR (left) and IF (right) analyses of LINE-1 mRNA and ORF1p protein, respectively, after transfection with LINE-1-specific (indicated as L1, map in Fig. 1 A) and luciferase-specific (GL2, control) siRNAs in PC3 cells. Histograms represent the relative expression of LINE-1 mRNA in control (GL2, taken as 1) and L1-interfered cells after normalization to GAPDH. Ratios were calculated using the 2-ΔΔCT method and p-values by the unpaired *t*-test. ***, p<0.001 (n, 8). The ORF1p signal intensity was measured by quantitative IF at the single cell level, selecting whole cells as the Region of Interest (RoI) (details in Methods). Graphs represent the distribution of ORF1p signal intensities (in arbitrary units) in individual cells (120 to 180 cells per condition were counted in 2 independent experiments), and statistically analyzed using the Mann-Whitney test: ****, p<0.0001. **C.** siRNA-mediated LINE-1 down-regulation prevents SPV122.2-dependent lamin B1 fragmentation. The upper panel shows representative confocal images of lamin B1 in DMSO-treated controls, either non-interfered (indicated as -, panel a), or interfered with neutral siRNA (+GL2, panel b), or with LINE-1-specific siRNA (+L1, panel c). In the lower panel lamin B1 is immunostained in SPV122.2-treated cultures either non-interfered (d), or interfered with GL2 (e), or interfered with LINE-1-specific siRNAs (L1, f). Sites of lamin B1 discontinuity are indicated by white arrows. The graph quantifies the fraction of cells exhibiting discontinuous lamin B1 (mean % ± SD values) from >100 cells/condition in 3 assays. Samples were statistically analyzed using the unpaired *t*-test. *, p <0.05. Bars, 20 μm.

Given that both SPV1122.2 and ABC increase full-length LINE-1 expression (Fig. 1A) and that both promote the formation of PLA ligation products between ORF1p and lamin B1, we assessed whether LINE-1 expression was also implicated in the induction of lamina disruption associated with SPV122.2 treatment. To that aim PC3 cells were transfected with LINE-1-specific small interfering (si) RNAs (depicted in the map in Fig. 1A, collectively indicated as L1 siRNA in Fig. 9B-C) or with control siRNA, directed against the firefly luciferase gene, absent in mammalian cells (indicated as GL2 siRNA in Fig. 9B-C). PC3 cell treatment with L1-specific siRNAs yielded reduced LINE-1 mRNA levels by at least ∼60%, resulting in a highly significant decrease in ORF1p protein abundance (Fig. 9B). PC3 cells interfered with either L1-specific or with control GL2 siRNAs were subsequently exposed to SPV122.2 and stained for lamin B1. In the absence of SPV122.2, cells displayed an intact lamin morphology, regardless of whether they were non-transfected or transfected with GL2 or L1 siRNA (Fig. 9C, a-c panels). Adding SPV122.2 to PC3 cultures induced lamin B1 damage in both untransfected (indicated as -, Fig. 9C, panel d) and GL2 siRNA-transfected cells (Fig. 9C, panel e), as expected; white arrows point to micronuclei and to lamin B1 fragments resulting from the nuclear envelope disruption. In contrast, siRNA-mediated reduction of LINE-1 expression yielded a significant reduction in cells exhibiting lamina damage caused by SPV122.2 (Fig. 9C, panel f). Thus LINE-1 silencing results in a protective effect over lamin B1, which suggests that LINE-1 expression has an active role in the lamin-disruptive effect of SPV122.2.

## Discussion

In earlier work we found that two NNRTIs, EFV and SPV122.2, both designed to target the HIV-encoded RT, reduce the proliferation capacity of HIV non-infected cancer cells, both in culture and in animal models^28, 29, 38, 51, 54^. Here we report that both EFV and SPV122.2, and the NRTI ABC, increase the expression of full-length L1 mRNA in PC3 prostate cancer cells. This is reflected by an increase in ORF1p protein abundance, particularly in nuclei. That increase activates a cascade of events, ultimately leading to autophagy-dependent nuclear rearrangements that are independent on genomic retrotransposition events. This finding was unexpected. It is well-established that increased L1 expression is a widespread response triggered by stressors and environmental stimuli^80–84^. This similarity suggests that the RTIs tested here act as stressors in cancer cells, which is consistent with their ability to induce DNA damage, as shown in this and in previous studies^54, 55, 85^.

The present finding that both ABC and SPV122.2 increase ORF1p abundance, particularly in nuclei in PC3 prostate cancer cells is consistent with results obtained in a distant biological context, i.e. a neuronal cell model exposed to oxidative stress, in which L1 mRNA and ORF1p were also up- regulated and accompanied by increased ORF1p translocation in the nucleus^86^. Interestingly, in that model, ORF1p established stress-induced interactions with nuclear proteins, including lamin B1, NUP153, a nucleoporin on the nuclear side of the nuclear pore, and importin beta, the major nuclear import receptor. The increase in L1 mRNA levels, the surge in accumulation of ORF1p in the nucleus, and the appearance of ligation products between ORF1p and nuclear proteins in such distant cellular contexts as PC3 cancer cells and neurons exposed to diversified stressors suggest a widespread ability of ORF1p to orchestrate the response to stress.

Key steps of the cellular response identified in this work are summarized in the model in Fig. 10. Treatment of PC3 cells with NNRTIs and NRTIs induced an increased accumulation of ORF1p in nuclei and the establishment of novel associations between ORF1p and nuclear factors that otherwise occur rarely and mostly at non-specific locations in control cells. RTI treatment causes ORF1p to localize at sites of genomic DNA damage, forming PLA products with γ-H2AX, which also increased in SPV122.2- and ABC-treated cells in consequence of the induction of DNA damage. The increase of γ−H2AX/ORF1p ligation products in RTI-treated cells suggests that impaired, interrupted or unfinished L1-retrotransposition events result in unrepaired DNA breaks generated by the L1 ORF2p-encoded EN activity. The γ−H2AX interactor BCLAF1 also increases and becomes another ORF1p-interacting partner in RTI-treated cells. A complementary MS-based analysis demonstrated that two other NRTIs, islatravir and stampidine, also increased the amount of γ-H2AX that co-immunoprecipitated with ORF1p. These findings are both consistent with and complementary to the PLA results. Together the data suggest that the recruitment of ORF1p to DSBs, and ORF1p’s ability to associate with DNA repair factors at DSBs, contribute to signaling from DNA damaged sites to the cellular response networks.

**Fig. 10.**
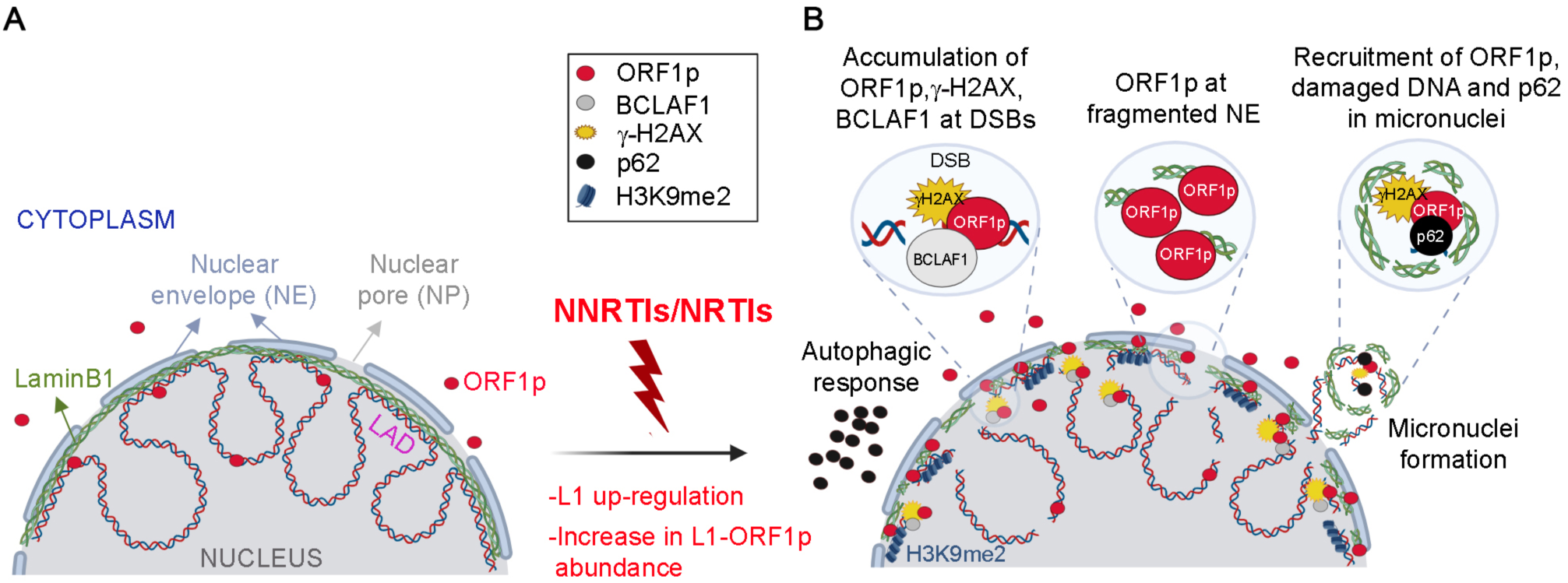
Proposed model for LINE-1-mediated nuclear remodeling induced by NNRTIs/NRTIs. **A.** Schematic of genome–lamin interactions in untreated cells. Genomic DNA (double strand blue/red) forms transcriptionally active chromatin loops across the nucleus and lamin-associated domains (LADs) of chromatin that are anchored to the nuclear envelope (NE) via lamin B1 (green). NNRTIs and NRTIs (red flash) up-regulate *L1* mRNA and ORF1p. **B.** L1 upregulation triggers an autophagic response substantiated by an increase of p62 (black dots) and LC3-II (not depicted), and a cascade of events: peripheral increase of heterochromatin marks (H3K9me2, blue), DNA damage (double strand breaks, DSBs), and accumulation of ORF1p. At DSBs ORF1p interacts with both γ−H2AX and BCLAF1; concomitant with this, lamin B1 fragments and ORF1p accumulates at sites of ruptures; lamin B1-coated micronuclei appear, which contain damaged DNA (γ−H2AX), ORF1p and p62.

It is well-established that DNA damage and nuclear envelope integrity are connected in a mutual interplay: on the one hand, lamin B1 disruption increases the susceptibility of genomic DNA to the induction of damage and activates the autophagic response^74, 87,88^; on the other hand, the DNA damage response network contributes in turn to induce ruptures in the nuclear envelope^73^. The nuclear lamina provides anchorage sites for chromatin domains. Therein, lamina-associated domains (LADs) are dynamic structures predominantly constituted by LINE-1 DNA^89^ (reviewed in^90,91^). We now show that both SPV122.2 and ABC, concomitant with inducing DNA damage and accumulating ORF1p at DSBs, also promote the accumulation of ORF1p at sites where the lamin B1 layer undergoes fragmentation and the formation of ligation products between ORF1p and lamin B1. BCLAF1, in addition to interacting with γ−H2AX and ORF1p at DSBs, is also reported to interact with emerin^92^, an inner nuclear membrane protein lying in close contact with lamin B1 at more than 1300 domains in the genome of human fibroblasts^93^. These data add support to the idea that ORF1p acts as part of a signaling mechanism connecting the damaged DNA and the nuclear envelope response.

In parallel, p62 also accumulates at sites of nuclear lamina ruptures induced by both SPV122.2 and ABC. Micronuclei are generated, which concentrate large amounts of ORF1p, γ−H2AX, and p62. Micronuclei formed under aneuploidy-inducing treatments also contain damaged DNA and recruit p62^78^; importantly, the latter inhibits the activity of the ESCRTIII complex, which operates in cellular trafficking and membrane budding and is required for nuclear envelope sealing, The accumulation of p62 and the ensuing inhibiton of ESCRTIII activate the autophagic response. Our experiments indicate that the micronuclei formed in cells treated RTIs are endowed with key components potentially capable of activating autophagy and ensuring the clearance of micronuclei containing the damaged DNA. Remarkably, the ESCRTIII complex can interact with L1RNPs and is reported to be critical for LINE-1 retrotransposition, suggesting a role for ESCRTIII in LINE-1 propagation^94^. It is also worth noting that LINE-1 mRNA, when overexpressed, triggers autophagy, which promotes the degradation of excess LINE-1 mRNA in response in order to prevent new retrotransposon insertions^95^.

To summarize, the present data show that RTIs, as many stressing agents, up-regulate the abundance of LINE-1 mRNA and ORF1p, induce ORF1p-centered interactions with DNA damage sensors and with the fragmented nuclear envelope via lamin B1, favor the formation of micronuclei, and activate autophagy, as indicated by the increased expression of autophagic markers LC3-II and p62. We show that the activation of autophagy is not only concomitant with the induction of DNA damage, lamina fragmentation, and expulsion of DNA-associated lamin B1 fragments in response to RTIs, but that it is actually required in order to elicit these events. Indeed, 3-MA, an inhibitor of the PI3 kinase that blocks autophagosome formation, suppresses both the recruitment of γ-H2AX accumulation at damaged DNA and nuclear lamina fragmentation. We further demonstrate that LINE-1 expression is required to elicit these effects, as siRNA-mediated reduction of LINE-1 expression significantly mitigates the induction of nuclear lamina fragmentation by RTIs. These events may represent functional steps in the propagation of a LINE-1-mediated response to stress throughout the genome. We speculate that cancer cells exposed to RTI-induced stress respond by increasing full-length LINE-1 expression, possibly in an attempt to increase their genome plasticity. That event can then initiate a retrotransposition-independent signal capable of activating the autophagic pathway in order to enable the removal of damaged subcellular structures and facilitating either an adaptive cell survival or the self-digestion and death of heavily damaged cells.

The finding that the treatment of PC3 cancer cells with all tested RTIs results in an increase in LINE-1 mRNA expression (a hallmark of cancer cells), while concomitantly exerting antiproliferative and anticancer effects, may appear paradoxical. It is worth noting, however, that increased expression of retroelements beyond a critical threshold has been reported to improve the cellular response to cancer therapy (reviewed in^27,96^). The threshold concept refers to the level of intermediate nucleic acid products, including retrotransposon-encoded double-stranded RNA, bidirectional transcription products, and cDNAs resulting from L1-mediated reverse transcription. These intermediates can be generated in the course of the retrotransposition cycle, are detected by cellular sensors, and are thought to become toxic for cancer cells when exceeding a threshold of tolerance. The results reported here attribute a central role to LINE-1 expression and ORF1p in modulating the cancer cell fate in response to drug-induced stress. Our experiments, designed to understand the capacity of RTIs to inhibit cancer cell proliferation, led to classifying them as stressors and hinted at the activation of a response pathway that ultimately reorganizes the nucleus and conveys the cells to death in an autophagy and LINE-1-dependent manner: thus, a role of the LINE- 1/autophagy connection is now beginning to emerge in this reorganization.

## Materials and Methods

### Cell cultures and drug treatments

The prostate carcinoma cell line PC3 (ATCC CRL-1435) was cultured in RPMI 1640 supplemented with 10% FBS, 1% L-glutamine and 1% penicillin/streptomycin in a humidified atmosphere at 37°C in 5% CO_2_. Five h after seeding cells were incubated with RTIs Efavirenz (EFV, 20 μM), SPV122.2 (20 μM) and Abacavir (ABC, 50 μM). EFV was purified from commercially available Sustiva (Roche). SPV122.2 stereoisomer was isolated from the parent racemic mixture by repetitive semipreparative chiral HPLC as described^97^. Both drugs were dissolved in DMSO (0.2% final concentration). PC3 cells cultured in 0.2% DMSO alone served as controls. ABC was purified from commercially available Ziagen tablets (GlaxoSmithKline) as described^52^ and dissolved in RPMI medium. Untreated PC3 cells were used as controls. All cells were harvested after 96 h of treatment (cycle 1), replated at the same initial density for another 96 h (cycle 2) and finally harvested for analysis at the end of cycle 2. Freshly prepared 3-Methyladenine (3-MA, Sigma-Aldrich M9281) was used 3 mM (final concentration) and added in the last 72 h of each cycle. Culturing conditions for N2102Ep cells were previously described^66^. For treatment with NRTIs, the cells were exposed to 10 μM Islatravir or Stampidine and harvested 72 h after treatment. Both compounds were dissolved in DMSO (0.1% [v/v] final concentration). Control cultures received DMSO alone at the same concentration.

### Western blot analysis

PC3 cells were seeded in a T-75 flask dish, collected by scrapping in ice-cold PBS with PMSF, centrifuged at 1,000 × g for 5 min at 4°C and washed twice with ice-cold PBS. Cells were lysed in M-PER (mammalian protein extraction reagent) according to manufacturer’s instruction (Thermo Scientific) plus protease inhibitor cocktail (Roche Life Science) and centrifuged at 12000 rpm for 20 min, 4°C. The protein concentration of centrifugally clarified extracts was determined using the Bradford Reagent (Sigma-Aldrich). 25-30 μg total cell lysate per lane were separated through 10- 12% SDS-polyacrylamide gels and transferred to nitrocellulose membranes (Thermo Scientific). The membranes were blocked in 1x TBS containing 0.1% Tween-20 and 5% BSA, then incubated (4°C, overnight) with primary antibodies against ORF1p, β-tubulin, GAPDH as indicated in Table S2. After washing, membranes were again incubated using secondary HRP-conjugated species-specific (anti-goat, ab205723, or anti-rabbit, ab205718) antibodies from Abcam. Images for individual blots were developed using the Clarity Western ECL Substrate (Bio-Rad #1705061) in the ChemiDoc XRS+ detection system (Bio-Rad). Endogenous Tubulin or GAPDH were used as a loading control and the signal intensities for each blot were quantified by calculating the band intensity of the protein of interest relative to either tubulin or GAPDH, as indicated, by densitometry using Image Lab software (Bio-Rad).

### Immunofluorescence (IF) analysis and confocal microscopy (CLSM)

For microscopy analyses PC3 cells were grown on sterile coverslips. IF staining of ORF1p, γ−H2AX, lamin B1, LC3-II and p62 were carried out on methanol-fixed cells (10 min, 4°C); H3K9me2 IF staining was performed in cells fixed in 4% PFA/30mM sucrose (10 min, room T°), permeabilized in 0.1% Triton X-100 (5 min, room T°) and incubated with glycine 0,1M (10 min, room T°). Fixed cells were blocked in 3% BSA, 0.05% Tween-20 in PBS (1 h, 37 °C) and incubated overnight at 4°C with primary antibody at the dilution indicated in Table S2. Rabbit primary antibodies were detected using anti-rabbit IgG secondary antibodies conjugated with either Cy3 (Jackson Immunoresearch 711- 165-152), or FITC (Jackson Immunoresearch 711- 095-152). FITC-conjugated goat anti-mouse IgG/IgM (Jackson Immunoresearch 155-095-068) and CY3.5-conjugated horse anti-mouse (Vector) were used for detection of mouse primary antibody. In all samples the DNA was counterstained with 0.1 μg/ml 4,6-diamidino-2-phenyl-indole (DAPI) and mounted in Vectashield (Vector Laboratories). Images were acquired under a Nikon Eclipse 90i microscope equipped with a Quicam Fast 1394 CCD camera (Qimaging) using either a 100× immersion oil objective (NA 1.3) for single-cell quantitative analysis, or a 40× objective (NA 0.75) for field-of-view analysis. For micronuclei analysis images were acquired under a Nikon Ti2 confocal spinning disk microscope implemented with a Crest X-Light V3 module (CrestOptics SpA) equipped with the Kinetix sCMOS camera (Teledyne Photometrics), a 60× (immersion oil, NA 1.4) objective and CELESTA lasers (Lumencor). Image projections from z-stacks were created using the Maximum Intensity Projection (MIP) function. Images were taken using the Nis-Elements AR 3.2 software; Nis-Elements AR 5.11 was used to quantify IF and PLA signals. Confocal images were acquired under a Leica TCS SP5 confocal microscope as described^54^ and analyzed using Image J processing software.

#### Proximity ligation assay (PLA)

Cells grown on coverslips were fixed and blocked as for IF, then processed for PLA using the Duolink® In Situ Detection Reagents orange DUO92007 (Sigma-Aldrich). Cells were fixed, blocked, and incubated with primary antibodies as for IF. Species-specific MINUS and PLUS PLA probes (DUO92004 and DUO92002, Sigma-Aldrich; 1:5 dilution in PBS containing 0.05% Tween-20 and 3% BSA) were then added (60 min, 37 °C, pre-heated humidity chamber). Ligation (30 min, 37°C) and amplification at 37°C for 40-70 min, depending on protein pairs^98^, were performed following the Olink Bioscience protocol. Cells were co-stained using DAPI and chicken β-tubulin antibody (Abcam ab41489), followed by FITC-conjugated anti-chicken IgG (Jackson Immunoresearch). Signals were quantified using the NIS-Elements 4.0/4.1 software (Nikon), either in whole cells using tubulin staining to delimit the cell profile, or in nuclei as the regions of interest (RoI), selected using DAPI.

#### Co-immunoprecipitation and Mass spectrometry

Cryomilled N2102Ep cell powder was used to affinity purify LINE-1 RNPs, as described^66^. Dynabeads M270 epoxy (Invitrogen) coupled with anti-LINE-1 ORF1p (clone 4H1) were used for immunoprecipitation. Protein extraction and co-immunoprecipitation were performed in 20 mM HEPES-Na (pH 7.4), 50 mM MgCl, and 0.1% (v/v) Tween 20. For the shotgun proteomics, the IP elutions were processed using S-Trap columns^99^ as described^66^. The raw files were collected using DDA mode injecting 5 µl of the final samples onto a 75 µm x 50 cm Acclaim PepMap RSLC nano Viper column filled with 2 µm C18 particles via a Dionex Ultimate 3000 HPLC system interfaced with an Orbitrap Exploris 480 mass spectrometer. Column temperature was set to 40°C. Using a flow rate of 300 nl/min, peptides were eluted in a gradient of increasing acetronitrile, where solvent A is 0.1% formic acid in water and solvent B is 0.1% formic acid in acetronitrile. Peptides are ionized by electrospray at 1.8 kV as they elute. The elution gradient is 60 min as follows: 3% B over 3 min; 3 to 50% B over 45 min; 2 min to 80% B; then wash at 80% B over 5 min, 80 to 3% B over 2 min and then the column was equilibrated with 3% B for 3 min. Full scans were acquired in profile mode at 120000 resolution. The top 25 most intense ions with charge state +2-6 in each full scan are fragmented by higher energy collisional dissociation.

#### Data processing

The computational proteomic pipeline used was previously described^100^. Peptide identification and quantification using MaxQuant (version 1.9) and searched against a human-specific proteomic database. The following setting was used: include contaminants - true; PSM & protein FDR -0.01; quantify unmodified peptides and oxidation (M), acetyl (protein N-term), methylthio (C); iBAQ-true; iBAQ log fit - false; match between runs - true; decoy mode – revert; advanced ratios - false; second peptides - true; advanced ratios - false; second peptides - true; stabilize large LFQ ratios - true; separate LFQ in parameter groups - true; razor protein FDR – true. Proteins marked as “contaminants” or “reverse” were removed. Only proteins which had “Peptide counts (razor + unique)” ≥ 2 were retained for analysis. Using the data obtained, protein intensities were log2 - transformed in order to do the imputation as described. For statistical testing, log2- transformed intensities were used for t-test two samples unpaired between ORF1p IP RTi Treatment and DMSO. Protein enriched with the ORF1p target in each experiment was considered statistically significant if the Benjamini-Hochberg adjusted p-value was ≤ 0.05 and log2 transformed fold-change (log2FC; treatment/control) was ≥ 1.

#### Statistical analysis of imaging data

IF intensity and PLA data were analyzed using GraphPad Prism 8. The following statistical tests were employed: Student’s *t* test to compare mean +/- SD values between independent groups; Mann-Whitney non-parametric test, or two-way ANOVA Dunn’s multiple comparisons test, to compare continuous values, which in most experiments had a non-normal distribution across samples between two or more independent groups, respectively. The distribution of values measured in single cells is represented in dot plots. All graphs show intervals of statistical significance and *p* values are indicated in the figure legends.

#### RNA isolation and cDNA synthesis

RTI-treated PC3 cultures and their respective controls were washed twice with cold PBS and lysed by direct TRIZOL addition to culture flasks and total RNA was immediately isolated according to the manufacturer’s instructions (Invitrogen, Carlsbad, CA). RNA integrity was confirmed on 1% agarose gels in TAE (1X). For each sample, 2 µg of total RNA was pretreated with DNAse I Amp Grade (Invitrogen, cat. 18068015) following protocol’s instructions. First strand cDNA synthesis, using oligo (dT)_20_, was performed in 20µl of final volume using SuperScript III Reverse Transcriptase (Invitrogen, cat. 18080-093).

#### RNA interference

Three 21-nt double-stranded siRNA oligonucleotides were designed to target the mRNA encoded by the L1.3 reference sequence (GenBank: L19088.1) in L1 ORF1 (Fig. 1A; ID s552967, 5’- AGAAGAAUGUAUAACUAGAtt-3’ [Pos. 1128-1146]; ID s552968, 5’- GUGGAUCUCUCGGCAGAAAtt-3’ [Pos. 1660-1678]) and in L1 ORF2 (Fig. 1A; ID s552969, 5’-GGUGAACUCCCAUUCGUAAtt-3’ [Pos. 4252-4270]). All L1-specific siRNAs were synthesized by Ambion^®^/Thermofisher. Briefly, 1.5×10^5^ cells were plated in a six-well dish (35 mm per well) in complete medium and incubated overnight at 37°C in a humidified 5% CO2 incubator. The following day, the cells were washed once with RPMI medium, then transfected using Lipofectamine RNAiMAX (Invitrogen) according to the manufacturer’s instructions. Two transfection mixtures were prepared for each experiment: the first mixture contained 40 nM of each L1-specific siRNA in RMPI medium, while the second mix had Lipofectamine RNAiMAX dissolved in RMPI. After combining the mixtures and incubating at RT for 25 min, the combined mixture was pipetted drop-wise into each well. The plates were incubated at 37°C in a humidified atmosphere of 5% CO_2_ for 6 h. The transfection medium was then removed and replaced with fresh complete medium. After 48 h, a second round of transfection with siRNA was performed and cells were incubated for an additional 24h. For control, cells were transfected with a double-stranded small RNA called GL2 (5′-CGUACGCGGAAUACUUCGATT-3′; Ambion^®^/Thermofisher) directed against the firefly luciferase, which is not expressed in mammalian cells.

#### Quantitative Real time PCR

Quantitative Real time PCR (RT-PCR) was performed in an IQ5 Real-Time PCR System (BIO-RAD) using PowerUp SYBR Green Master Mix (Applied Biosystems, cat. A25741) under the following conditions: one cycle at 50°C for 2 min, one cycle at 95°C for 2 min, 40 cycles at 95°C for 10 s, 60°C for 30 s. The relative expression of L1ORF1 and L1ORF2 was determined by the 2^ΔΔ^Ct method, normalized on GAPDH expression chosen as housekeeping control. Samples from three independent experiments were analyzed by qPCR and each sample was routinely analyzed in triplicate. All data were expressed as mean fold change ± standard deviation (SD).

The following primers were used for Real Time PCR are L1ORF1 For: CCGATGCGATCAACTGGAAG (Pos. 1236-1255), L1ORF1 Rev: TGCCTTGCTAGATTGGGGAA (Pos. 1484-1465), L1ORF2 For: TGAAAACCGGCACAAGACAG (Pos. 3947-3966), L1ORF2 Rev: GCTGAGACGATGGGGTTTTC (Pos. 4133-4114), BCLAF1 For: CCATCCCCTTGGTATTTGTC, BCLAF1 Rev: GGCCAAAAGAGGAAGAGGTC, GAPDH For: CCC TTC ATT GAC CTC AAC TAC ATG, GAPDH Rev: TGG GAT TTC CAT TGA TGA CAA GC. Nucleotide positions of L1-specific primers refer to the L1.3 reference sequence (GenBank: L19088.1). Differences in L1/GAPDH ratios were measured between NNRTIS (EFV and SPV122.2) and DMSO, and between ABC and untreated controls, and statistically analyzed using the two-way ANOVA with Dunn’s post-test multiple comparisons.

## Acknowledgements

We are grateful to Dr. Vincenzo Costanzo for experimental contributions to imaging experiments in this work. We thank Shaoshuai Xie for assistance with RTI-treated cell processing and Luciano Di Stefano for MS data collection. Stampidine and islatrivir used in the IP-MS experiments were kindly provided by Rome Therapeutics and these experiments were funded in-part by the PPP allowance made available by Health∼Holland, Top Sector Life Sciences & Health, to stimulate public-private partnerships.

## Funding

This work was supported by funds from “Fondazione Roma”, project “Investigating the cellular endogenous Reverse Transcriptase as a novel therapeutic target and an early tumor marker” (to CS); IFT-CNR project DSB.AD007.088 (to AS); MUR-PRIN grant 2017FNZRN3_005 and MUR PON-IR program for research infrastructures PIR01_00023 “IMPARA - Imaging from molecules to the preclinics” (to PL); AIRC IG 2020 (grant 24942) (to DT); National Institutes of Health (NIH) grants R01AG078925 and R01AI186337 (to JL).

## Conflicts of interest

The authors declare no conflict of interest. JL reports personal fees and equity from Rome Therapeutics, for consulting services provided outside the submitted work.

## Author contributions

M. Damizia: methodology, investigation, formal analysis, validation; M. Baranzini: methodology, investigation, formal analysis, validation; I. Sciamanna: conceptualization, methodology, investigation, validation; L. Altieri: methodology, investigation, validation; P. Rovella: methodology, investigation, validation; O.G. Rosas Bringa: methodology, formal analysis, software; S. Hozeifi: methodology, investigation; L.J. Saba: investigation, validation; A. Mourtzinos: investigation, validation; A. O’Shea: investigation, validation; M.S. Zeya Ansari: investigation, validation; F. Andreola: investigation, validation; R. Cirilli: resources; writing - review & editing; G. Sbardella: resources; writing - review & editing; A.L. Serafino: methodology, investigation, formal analysis, writing - review & editing, funding acquisition; G.G. Schumann: conceptualization, methodology, writing - review & editing; D. Trisciuoglio: conceptualization, methodology, investigation, supervision, writing, funding acquisition; J. La Cava: methodology, supervision, writing, funding acquisition; P. Lavia: conceptualization, methodology, supervision, writing, funding acquisition; C. Spadafora: conceptualization, methodology, supervision, writing, funding acquisition.

## Data availability

All relevant data are within the manuscript and its Supplementary files. The data are available from the lead authors on reasonable request.

## Notes

### Competing Interest Statement

The authors declare no conflict of interest. J. Lacava reports personal fees and equity from Rome Therapeutics, for consulting services provided outside the submitted work.

### Summary of Updates

This revised version is complete and contains 10 figures. In the originally submitted version, Figure 7 was inadvertently not loaded with the manuscript

