## supplementary text for "Reverse transcriptase inhibitors induce autophagy in a LINE-1 ORF1p-dependent manner"

**Short title: LINE-1 and autophagy**

**Keywords:** LINE-1, L1-ORF1p, RT inhibitors, γ−H2AX, lamin B1, micronuclei, autophagy, prostate cancer cells

**List of supplementary Files**

**Table S1.** Protein Enrichment in ORF1p IP on treatment with Stampidine vs DMSO and Islarivir vs DMSO

**Table S2.** Primary antibodies and conditions used in this study

**Fig. S1.** Both SPV122.2 and ABC up-regulate the abundance of autophagy markers in PC3 cells. **A.** Single-cell IF analysis of the autophagy markers LC3-II (green) and p62 (red): in SPV122.2-treated compared to DMSO control cultures. Upon treatment with SPV122.2, both proteins are up-regulated and redistribute to smaller cytoplasmic granules comparted to DMSO cultures. **B.** Single-cell IF analysis of LC3-II and p62 in ABC-treated cultures. ABC induces a somewhat weaker increase in the amount of both p62 and LC3-II proteins, as revealed by the signal intensity measured in fluorescence units (compare to SPV 122.2), yet statistically highly significant compared to the untreated control (CTR). Bars, 10 μm. The signal intensity distribution measured in single PC3 cells, treated with and without RTI, is displayed in the graphs below (30 to 40 cells per condition were measured in this experiment and reproduced in two assays, with 90 to 135 total counted cells per condition). The distributions were statistically analyzed using the Mann-Whitney test. **, p <0.0016; ***, p 0.0002; ****, p<0.0001

**Fig. S2**. SPV122.2 or ABC up-regulate the DNA damage and stress-responsive transcription factor BCLAF1 in PC3 cells. Representative IF fields from PC3 cultures treated either with SPV122.2 **(A)** or ABC **(B)**, stained for BCLAF1 (green). BCLAF1 signal intensity was measured on the DAPI mask (blue). Bars, 10 μm. The adjacent graphs display the BCLAF1 signal intensity/cell measured in RTI-treated cultures and in their respective controls (DMSO-treated or untreated, CTR) in 50 to 70 counted cells per condition. The results were reproduced in two assays, with 75 to 115 total analyzed cells per condition. The statistical analysis was performed using the Mann-Whitney test ****, p<0.0001.

**Fig. S3**. IF analysis depicts similar effects of SPV122.2 and ABC over lamin B1 and lysine 9-methylated histone H3 reorganization in PC3 cells. **A.** ABC (panel b) induces lamin B1 fragmentation (green), often associated with chromatin containing methylated histone H3 (depicted using anti-H3K9me2 antibody, red) expelled from the nucleus, suggestive of micronuclei formation. These structures are absent in PC3 untreated control cells (panel a). **B.** Similar structures are observed in SPV1222-treated PC3 cultures (panel b), but not in DMSO-treated (panel a) controls.
