## supplementary table 2 for "Reverse transcriptase inhibitors induce autophagy in a LINE-1 ORF1p-dependent manner"

**Supplementary Table 2. Primary antibodies used in this study**

| Antibody | Host | Source | Catalog | analysis | Dilution |
| --- | --- | --- | --- | --- | --- |
| BCLAF1 | rabbit | Thermo Fisher | PA5-55686 | IF, PLA | 1:500 |
| GAPDH | mouse | Santa Cruz | SC-47724 | WB | 1:1000 |
| γ-H2AX | rabbit | Cell signaling | 9718S | IF, PLA | 1:400 |
| H3K9met2 | mouse | Millipore | 07-411 | IF | 1:250 |
| Lamin B1 | rabbit | Abcam | Ab16048 | IF, PLA | 1:400 |
| Lamin B1 | chicken | Abcam | Ab90169 | IF | 1:100 |
| LC3B/LC3-II | rabbit | Cell Signaling | 2775S | IF | 1:250 |
| LC3B/LC3-II | rabbit | Sigma | L7543 | WB | 1:1000 |
| ORF1p (anti-L1) | mouse | Millipore | MABC1152 | IF, PLA, WB | 1:50/1:2000 |
| p62/SQSTHM1  α-tubulin | mouse  mouse | Santa Cruz  Sigma | SC-28359  T9026 | IF, WB  IF, WB | 1:250/1:200  1:1000 |
| β-tubulin | rabbit | Abcam | Ab6046 | IF | 1:300 |
| β-tubulin | chicken | Abcam | Ab41489 | PLA | 1:100 |
