## Supplementary figures and images for "Reverse transcriptase inhibitors induce autophagy in a LINE-1 ORF1p-dependent manner"

### supplementary fig.1

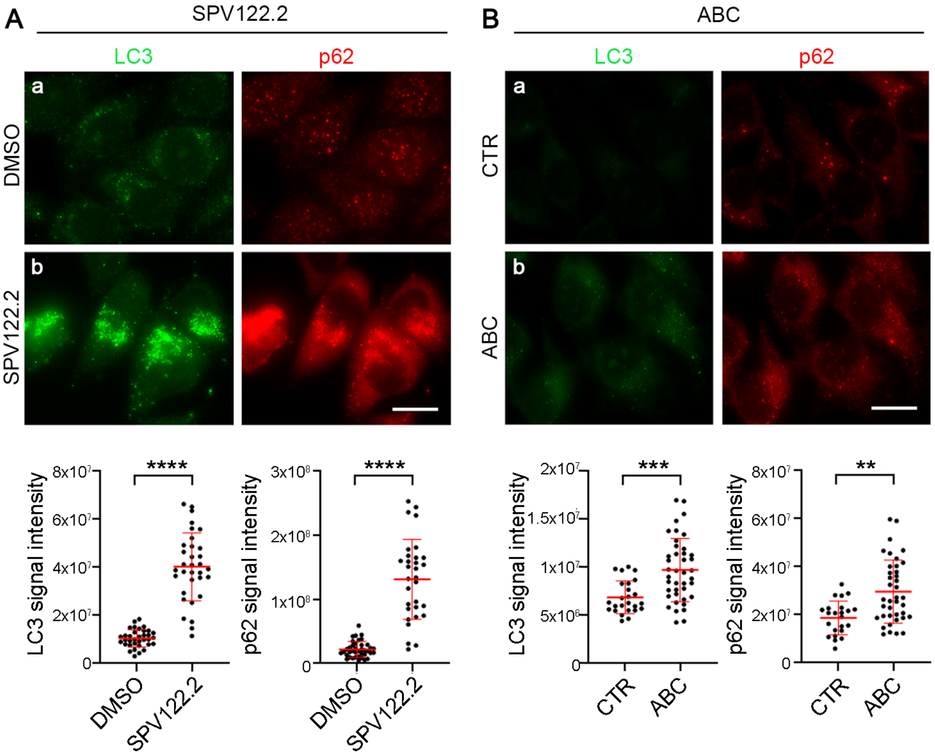

### supplementary fig.2

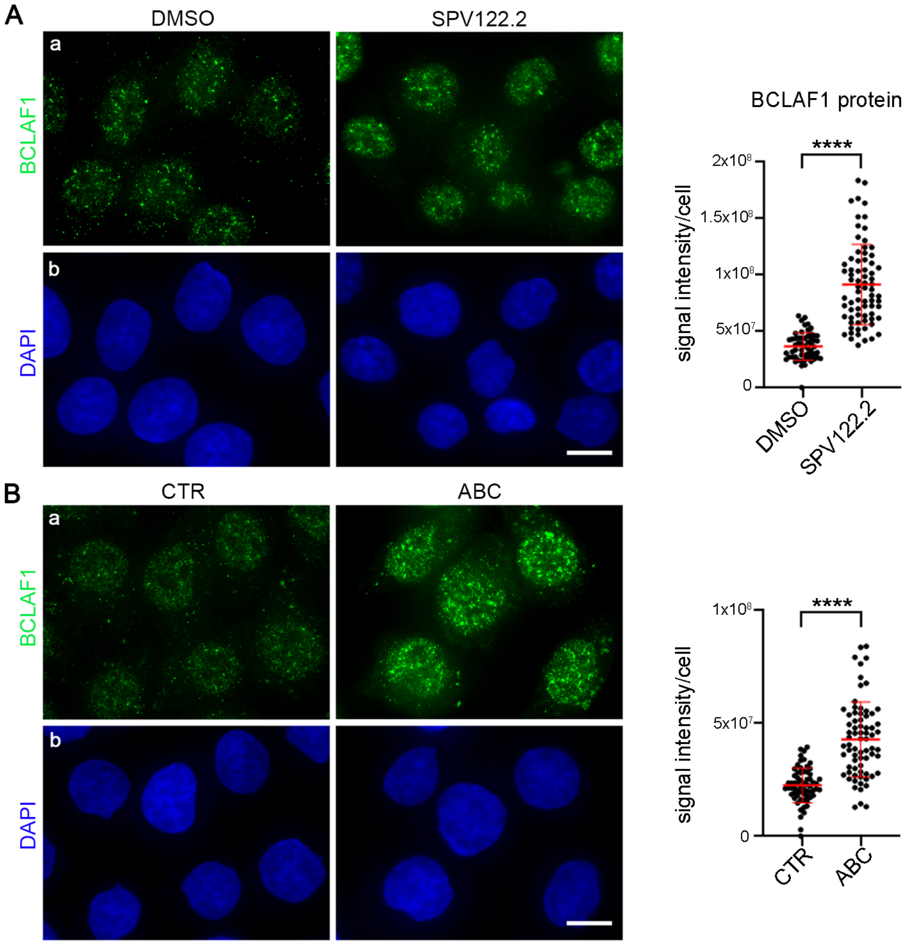

### supplementary fig.3

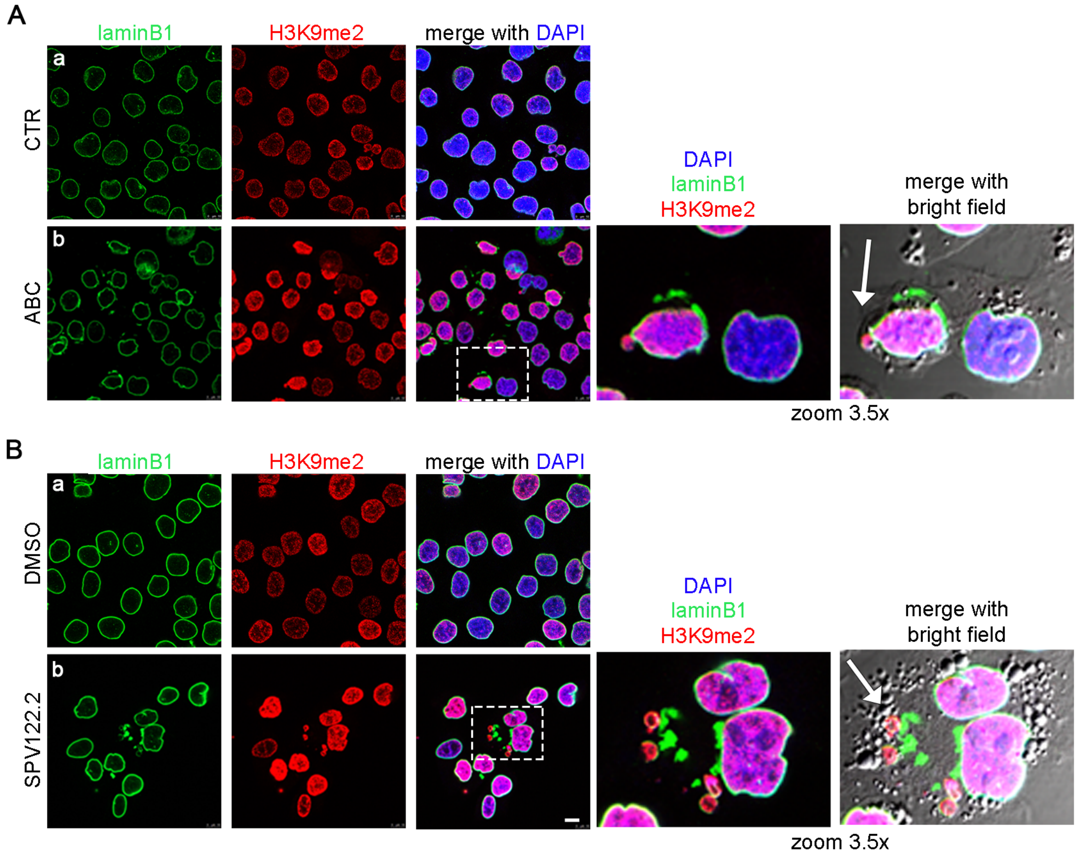
